# KMCP: accurate metagenomic profiling of both prokaryotic and viral populations by pseudo-mapping

**DOI:** 10.1101/2022.03.07.482835

**Authors:** Wei Shen, Hongyan Xiang, Tianquan Huang, Hui Tang, Mingli Peng, Dachuan Cai, Peng Hu, Hong Ren

**Affiliations:** Key Laboratory of Molecular Biology for Infectious Diseases (Ministry of Education), Institute for Viral Hepatitis, Department of Infectious Diseases, The Second Affiliated Hospital, Chongqing Medical University, Chongqing 400010, China

**Keywords:** Metagenomic profiling, *K*-mer, Genome chunks, Pseudo-mapping

## Abstract

**Motivation:** The growing number of microbial reference genomes enables the improvement of metagenomic profiling accuracy but also imposes greater requirements on the indexing efficiency, database size, and runtime of taxonomic profilers. Additionally, most profilers focus mainly on bacterial, archaeal, and fungal populations, while less attention is paid to viral communities.

**Results:** We present KMCP, a novel *k*-mer-based metagenomic profiling tool that utilizes genome coverage information by splitting the reference genomes into chunks and then stores *k*-mers in a modified and optimized COBS index for fast alignment-free sequence searching. KMCP combines *k*-mer similarity and genome coverage information to reduce the false positive rate of *k*-mer-based taxonomic classification and profiling methods. Benchmarking results based on simulated and real data demonstrate that KMCP, despite a longer running time than all other methods, not only allows the accurate taxonomic profiling of prokaryotic and viral populations but also provides confident pathogen detection in clinical samples of low depth.

**Availability and Implementation:** The software is open-source under the MIT license and available at https://github.com/shenwei356/kmcp.

## 1 Introduction

High-throughput sequencing technologies and computational methods have been combined in shotgun metagenomics approaches to study the composition and dynamics of microbial communities composed of bacteria, archaea, viruses, and fungi (Quince *et al*., 2017). Metagenomics is an essential means of exploring microbial diversity (Caporaso *et al*., 2012), the interaction between microorganisms and hosts (Human Microbiome Project, 2012), and the rapid detection of uncultivated and difficult-to-cultivate pathogens (Chiu and Miller, 2019). Metagenomic sequencing is mainly based on next-generation high-throughput sequencing platforms, which can generate millions of 50-250 bp short reads from each sample. Reference-based metagenomic profiling tools map these sequencing reads to a reference database and resolve the species present in the sample and their relative abundances (Sczyrba *et al*., 2017).

Several studies have assessed metagenomic profiling tools for taxonomic classification and profiling (Meyer *et al*., 2022; Sczyrba *et al*., 2017; Sun *et al*., 2021; Ye *et al*., 2019). Based on the methodology involved, these tools can be categorized into four types: (1) *k*-mer (length-*k* subsequences) based, such as Kraken2 (Wood *et al*., 2019), Bracken (Lu *et al*., 2017), Centrifuge (Kim *et al*., 2016) and Ganon (Piro *et al*., 2020); (2) alignment-based, such as DIAMOND (Buchfink *et al*., 2015), Kaiju (Menzel *et al*., 2016), DUDes (Piro *et al*., 2016), and SLIMM (Dadi *et al*., 2017); (3) marker-gene based, such as MetaPhlAn2 (Truong *et al*., 2015) and mOTUs2 (Milanese *et al*., 2019); and (4) machine learning-based, such as fastDNA (Menegaux and Vert, 2019), DeepMicrobes (Liang *et al*., 2020), CNN-RAI (Karagöz and Nalbantoglu, 2021), and BERTax (Mock *et al*., 2022). *K*-mer-based methods utilize exact *k*-mer matching, which is fast but sensitive to genomic mutation and variation because a single point mutation will result in at most *k* mismatches of *k*-mers. The taxonomic assignment algorithm based on the lowest common ancestor (LCA) often shows low accuracy at low taxonomic ranks because of closely related species sharing a large fraction of *k*-mers, and this accuracy decreases further with the increasing number of reference genomes (Nasko *et al*., 2018). In contrast, alignment-based methods show high tolerance for base variation. For example, SLIMM (Dadi *et al*., 2017) performs read mapping with Bowtie2 (Langmead and Salzberg, 2012) and maps matched reads into bins of fixed width across the reference genome; then, it utilizes the coverage information of reference genomes to remove unlikely genomes and obtain more uniquely mapped reads for assignment at lower ranks. Marker gene approaches perform well in the identification of archaea and bacteria (Meyer *et al*.,2022), while it is difficult to accurately identify viruses with this strategy since viruses do not have universally conserved genes, such as the 16S and 18S rRNA genes (Breitwieser *et al*., 2019).

The accuracy of metagenomic profiling also depends on the size and quality of the reference databases, and more extensive and comprehensive databases tend to yield better results (Breitwieser *et al*., 2019; Nasko *et al*., 2018; Piro *et al*., 2020). Databases such as NCBI RefSeq (O’Leary *et al*., 2016) and GenBank (Sayers *et al*., 2022) contain increasing numbers of microbial genomes (e.g., 250,000 assemblies belonging to 48,000 species in RefSeq release 210), which poses a major challenge to taxonomic profilers in terms of database size and construction efficiency. Recently, several genome catalogs have been created with the aim of providing comprehensive, high-quality reference genomes for taxonomic and functional research, such as human gut prokaryotic genome collections (UHGG (Almeida *et al*., 2021), HumGut (Hiseni *et al*.,2021), and an archaeal catalog (Chibani *et al*., 2022)) and human gut virus catalogs (GVD (Gregory *et al*., 2020), GPD (Camarillo-Guerrero *et al*., 2021), and MGV (Nayfach *et al*., 2021)). Sourmash (Irber *et al*., 2022) outperformed other tools in the Critical Assessment of Metagenome Interpretation (CAMI) based on mouse gut datasets (Meyer *et al*., 2019) using FracMinHash sketches with a scale of 10,000 calculated from all 141,677 prokaryotic genomes in a RefSeq snapshot. The sketching algorithm uses a small number of signatures from whole-genome sequences; therefore, it can index a large number of reference genomes. However, it is limited by reduced sensitivity in small genomes such as those of viruses.

Several computational approaches have been introduced to index and query large collections of sequencing datasets and genomes over the last few years (Marchet *et al*., 2021). Color-aggregative methods index the union set of *k*-mers, then associate information to each *k*-mer to its origin. For example, Bifrost (Holley and Melsted, 2020) and Cuttlefish 2 (Khan *et al*., 2022) use compacted de Bruijn graphs, and Mantis (Pandey *et al*., 2018) uses the counting quotient filter for *k*-mer indexing and exact querying. While *k*-mer aggregative methods index each dataset using a separate Bloom filter and then build an aggregation data structure for approximate querying. Sequence Bloom Tree (SBT) approaches (Harris and Medvedev, 2020; Solomon and Kingsford, 2016; Solomon and Kingsford, 2018; Sun *et al*., 2018) are designed for collections with high *k*-mer redundancy, such as human RNA-seq sequencing data. In contrast, Bloom filter matrix-based methods, including the BItsliced Genomic Signature Index (BIGSI) (Bradley *et al*.,2019), Compact Bit-Sliced Signature Index (COBS) (Bingmann *et al*., 2019), and Interleaved Bloom filter (IBF) (Dadi *et al*., 2018) approaches, are suitable for heterogeneous *k*-mer sets, such as those of microbial genomes. Instead of performing inexact pattern matching in alignment methods, *k*-mer-based methods simply examine the fraction of matched *k*-mers in the query sequence. Although the Bloom filter may return false positives due to its probabilistic data structure, the false-positive rate of a query sequence decreases exponentially with the number of query *k*-mers (Bingmann *et al*., 2019; Solomon and Kingsford, 2016). These features indicate that *k*-mer-based Bloom filter matrix methods can be applied to index reference genomes for taxonomic profiling. Among these methods, IBF has been used for read mapping in DREAM-Yara (Dadi *et al*., 2018) and read classification in Ganon (Piro *et al*., 2020).

The increasing number of reference genomes calls for new taxonomic profiling tools with high efficiency in database building and high accuracy in prokaryotic and viral populations. Here, we present a novel taxonomic profiling tool, KMCP (*K*-mer-based Metagenomic Classification and Profiling), that uses a modified COBS data structure to index reference genomes and combines *k*-mer similarity and genome coverage information to improve taxonomic profiling accuracy. We evaluate KMCP based on simulated and real metagenomic datasets composed of data from archaea, bacteria, and viruses, and the benchmarking results indicate that KMCP performs well in both prokaryotic and viral populations.

## 2 Methods

### 2.1 Overview

We have designed and implemented KMCP, a novel *k*-mer-based taxonomic profiling tool. KMCP provides modular subcommands for *k*-mer computation, index construction, read searching, search result merging, and taxonomic profiling (Supplementary Table S1 and Fig. S1). To reduce the high rate of false positives observed in traditional *k*-mer-based methods, we split the reference genomes into chunks (Fig. 1a) and then track both the origin and the approximate location (chunk number) in the reference for each query (Fig. 1b). In the profiling step, a three-round filtering step, with criteria including the *k*-mer similarity and the coverage information of genome chunks, is adopted to filter out suspicious matches for increasing profiling specificity. Then the taxonomic abundance is estimated using the Expectation-Maximization (EM) algorithm (Dempster *et al*., 1977) (Fig. 1c).

**Fig. 1.**
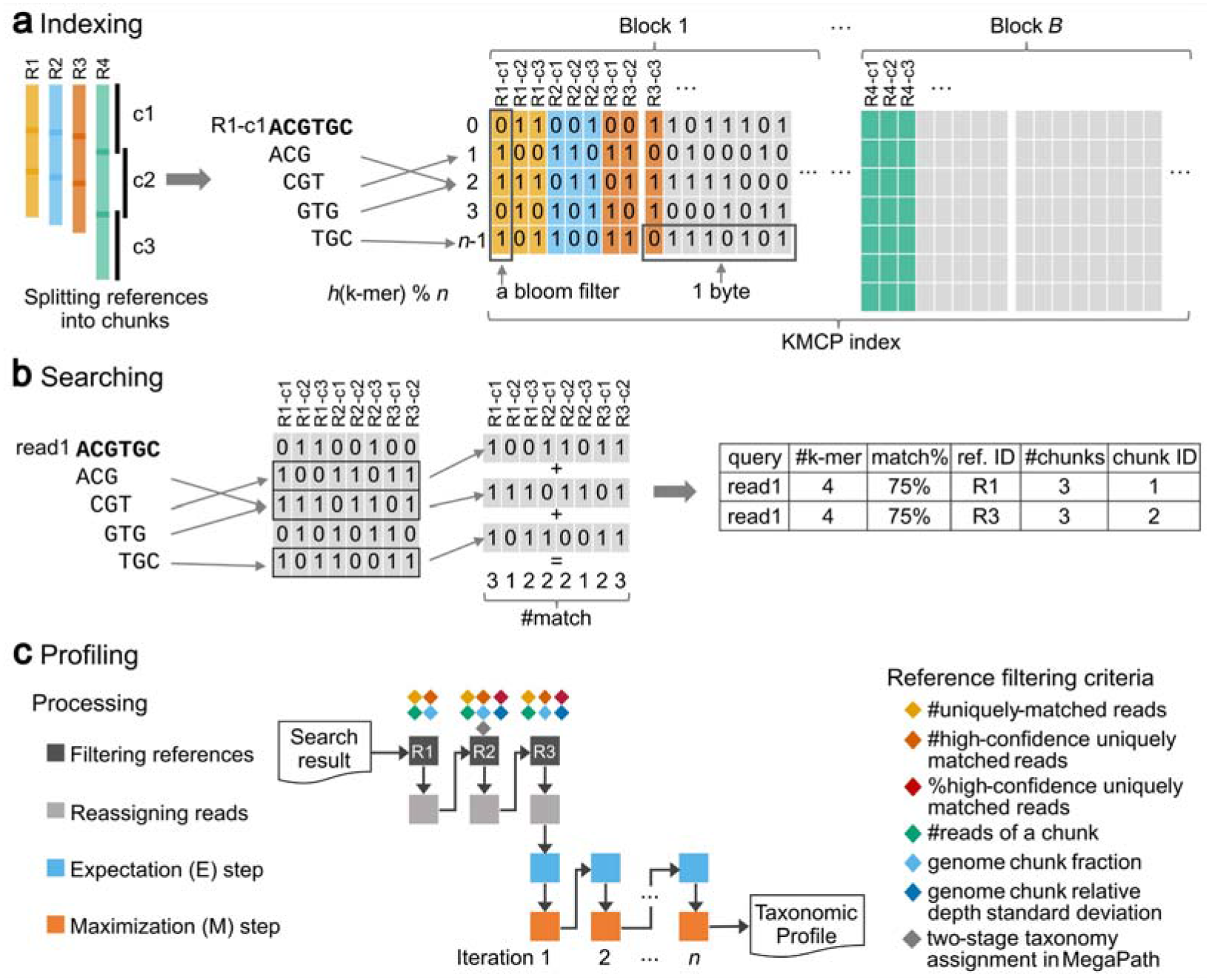
Schematic representation of KMCP. **(a)** Reference genomes are split into chunks, and the *k*-mers are stored in a modified COBS index. **(b)** The *k*-mers of a read are computed and searched in the index, and reference chunks sharing enough matched *k*-mers with the query are returned. **(c)** Matched references are filtered based on criteria including *k*-mer similarity and chunk coverage information, and relative abundances are estimated using the EM algorithm. Match% refers to the *k*-mer coverage (the proportion of matched *k*-mers and distinct *k*-mers of a query).

The read mapping process in KMCP is referred to as *pseudo-mapping*, which is similar to but different from the lightweight algorithm in Sailfish (Patro *et al*., 2014), pseudoalignment in Kallisto (Bray *et al*., 2016), quasi-mapping in RapMap (Srivastava *et al*., 2016), and lightweight mapping in Salmon (Patro *et al*., 2017). All these methods seek to elide the computation of base-to-base alignment while using distinct strategies (Srivastava *et al*., 2016). In KMCP, each reference genome is pre-split into chunks of equal size, and the *k*-mers of a query, as a whole, are compared to each genome chunk to find all possible ones sharing a predefined proportion of *k*-mers with the query. Like quasi-mapping in RapMap, KMCP tracks the target and position for each query. However, the read position in KMCP is approximate and in a predefined resolution (the number of genome chunks).

### 2.2 Indexing

KMCP efficiently builds a database from a collection of genome sequences using a modified COBS data structure (Fig. 1a). Plasmid sequences, which can be found in different genomes and affect later analysis, are filtered out according to sequence name. Then the microbial genomes are split into ten chunks with 150-bp overlaps by default, and the chunk information is further utilized in taxonomic profiling. The overlap size can be adjusted to the read length for single-end searching mode or the library fragment size for paired-end search mode. For genomes without a single complete genome sequence, chromosomes or contigs are concatenated with intervals of *k*-1 bases of N’s to avoid introducing fake *k*-mers. Unlike sketching-based methods like Sourmash, all nonredundant nucleotide canonical *k*-mers (*k*=21, by default) of each genome are hashed into 64-bit unsigned integers with ntHash (Mohamadi *et al*., 2016). Then, *k*-mer hashes alongside the genome chunk number and genome size are saved into binary files with the extension *.unik*. Next, these *k*-mer hashes are stored in a modified COBS index. Briefly, all *.unik* files are sorted in ascending order of the stored *k*-mer hash count and divided into blocks of *N* genome chunks. Then, *N* Bloom filters of equal size, with each Bloom filter corresponding to a genome chunk, are constructed and concatenated in a column-wise manner into a byte matrix for each block. Based on a given number of hash functions and a desired false-positive rate, the size of the Bloom filters is determined by the number of *k*-mers in the largest genome chunk. Other bloom filters will have the same or lower false-positive rate due to fewer k-mers (Supplementary Fig. S2). To avoid generating extremely large Bloom filters for the last block with the largest genome chunks, we set three thresholds (Supplementary Table S2) to create extra small indexes with fewer chunks, resulting in a smaller database size than COBS with the same parameters (Supplementary Table S3). In addition, the index files of all blocks are not concatenated to a large one for better parallelization of index loading and query searching. In summary, we have modified the COBS index structure to support genome chunk information, improved the indexing speed, and reduced the index size (Supplementary Table S3). A complete list of differences in algorithm and data structure between KMCP and COBS is listed in Supplementary Table S4.

Taxonomic information, including NCBI taxonomy dump (taxdump) files and a plain text file mapping reference identifiers to TaxIds, is only needed in the profiling step (Supplementary Fig. S1), allowing the database to be easily updated with newer taxonomy information. The taxdump files can be downloaded from the NCBI Taxonomy database (Schoch *et al*., 2020) or created from other taxonomy databases, including GTDB (Parks *et al*., 2022) and ICTV (Lefkowitz *et al*., 2018), using the *create-taxdump* command in TaxonKit (Shen and Ren, 2021). These index files and taxonomic information, including NCBI taxonomy dump files and TaxId mapping files, comprise the KMCP database.

### 2.3 Searching

For sequence searches against the KMCP database, *k*-mers of a single-end read or paired-end reads are computed with the parameters of the database and sent to query against Bloom filters of all index files in parallel. Although COBS shows a good query performance relative to similar tools, it is insufficient for shotgun metagenomic sequencing analysis (Supplementary Fig. S3). In our implementation, steps including index file loading, *k*-mer computation and hashing, and Bloom filter querying are highly optimized to take full advantage of modern multiple-core CPUs and high-speed solid-state disks (details are described in Supplementary Section 1.2). As a result, we have achieved a ten-fold increase in the speed of short-read batch searching relative to COBS (Supplementary Fig. S3).

Due to mutation or intraspecific polymorphism, a low *k*-mer similarity threshold is needed to identify querying sequences from distant homologous genomes during the search. At the same time, the Bloom-filter-based probabilistic index required enough matched *k*-mers to reduce the false-positive rate. To balance the index size, searching accuracy, and searching speed, we set the desired false-positive rate of the Bloom filters as 0.3 and use one hash function by default, following BIGSI and COBS. During the search step, the threshold of *k*-mer coverage (proportion of matched *k*-mers and unique *k*-mers of a query) is set to 0.55 (approximately equal to a sequence identity of 96.5%, Supplementary Fig. S4d), which limits the false-positive probability of a query to a reasonable level. Using multiple hash functions could reduce the false-positive rates of a single k-mer at the cost of searching speed; however, it is not necessary. The false-positive rate of 0.3 is high for a single k-mer; nevertheless, the value of a query sequence decreases exponentially with the number of query *k*-mers (Bingmann *et al*., 2019; Solomon and Kingsford, 2016), e.g., 5.30e-10 for 150-bp reads with 72 *k*-mers matched and 9.51e-07 for 100-bp reads with 44 *k*-mers matched, according to formula (4) in SBT article (Solomon and Kingsford, 2016) (Supplementary Fig. S5).

Unlike LCA-based methods, which retrieve the TaxId of each *k*-mer in the read and assign the LCA of the resulting TaxIds to the read, KMCP searches across the database with all *k*-mers and returns reference genome chunks sharing enough *k*-mers with the query. The searching step does not assign any taxonomic label to the query reads; therefore, search results from multiple databases can be merged. The mergeability of the search results makes it possible to construct separate databases for different reference genome datasets and choose various databases to search with considerable flexibility. Since the search speed is linearly related to the number of reference genome chunks in the databases (Supplementary Fig. S6b) and the index data (Bloom filters) of all genome chunks are independent, users can build smaller databases from partitions of the reference genomes and merge the results after searching against all small databases. The searching step can be parallelized with a computer cluster in which each computation node searches against a small database (Section 3.5). Computers with limited main memory can also utilize an extensive collection of reference genomes by building and searching against small databases.

### 2.4 Profiling

Due to the existence of homologous regions of genome sequences across multiple microbial species, methods solely relying on sequence similarity information suffer from a high false-positive rate. The genome coverage information can help determine the existence of a genome, thus reducing false positives (Dadi *et al*., 2017). Besides, a low sequence similarity threshold is essential for detecting microbes considering intraspecies variation, but it also brings false positives. Therefore, we require a matched reference genome having a minimum proportion (10% by default, Supplementary Table S5) of high-confidence uniquely matched reads with high *k*-mer coverage (≥0.75, approximately equal to ≥98% sequence similarity).

A three-round filtering step, with criteria including the *k*-mer similarity and genome chunks coverage information, is adopted to filter out suspicious matches (Fig. 1c). The first round adopts four relatively loose criteria: reference genomes with 1) at least one uniquely matched read and 2) at least one high-confidence uniquely matched read are collected and filtered by 3) the minimum fraction (0.8 by default) of genome chunks with 4) a minimum number (50 by default) of matched reads. If a query has multiple matches from different genomes of the same species, it is still considered a uniquely matched read. For the second round, all reads are reassigned, and the matched reference genomes are filtered with the above criteria but with different thresholds (20 for minimum uniquely matched reads and 5 for minimum high-confidence uniquely matched reads). And two additional criteria are added, including 5) the maximal standard deviation of relative genome chunk depths (2 by default) and 6) the minimum proportion of high-confidence uniquely matched reads (10% by default). Besides, the two-stage taxonomy assignment algorithm in MegaPath (Leung *et al*., 2020) is used to reduce further the spurious matched references. For the third round, all reads are reassigned, and the matched reference genomes are filtered with the same criteria as in round two, except for the two-stage taxonomy assignment algorithm. If all of the references explaining the matches of a read are filtered out, then the read is excluded from the next round.

Next, the relative abundances of all matched references are estimated using a standard EM algorithm (Fig. 1c). Initially, multi-mapped reads are assigned to matched references equally, and the taxonomic abundance of each reference is initialized with the cumulative matched bases and normalized by the genome size (the total bases of either complete genome or unfinished genomes like metagenome-assembled genomes with plasmid sequences filtered out). In each iteration of the EM algorithm, multi-mapped reads are reassigned based on the taxonomic abundance of matched references estimated in the previous iteration. The EM steps stop after a predefined iteration number (10 by default) or when a convergence criterion (the percentage change of the predominant genome is smaller than 0.01) is met.

We preset six profiling modes for multiple analysis scenarios, including mode 0 for pathogen detection, mode 1 for higher recall, mode 2 for high recall, mode 3 for the default mode, mode 4 for high precision, and mode 5 for higher precision (Supplementary Fig. S6d). The default values of the above filtering criteria for the six modes are listed in Supplementary Table S5. Users can choose a profiling mode and adjust parts of the parameters as required. For the pathogen detection mode, which needs to detect microbes from low-depth data, reference filtering criteria are adjusted to have the highest sensitivity. At least two of ten chunks are required to be covered with a minimum number of one read. In the extreme case, if a genome has only two reads matched in two genome chunks, each read having a length of 30 bp with 7 out of 10 *k*-mer matched. The false-positive rate for each read is 1.59e-03 (the threshold is 0.01), and the value for the matched genome is only 4.78e-07 according to Theorem 2 in the SBT article (Solomon and Kingsford, 2016), which applies to the matching of genome chunks too where chunks can be treated as repeated independent events in a model with a binomial distribution. Besides, an extra flag is switched on for only keeping main matches by abandoning matches with sharply decreased *k*-mer coverage, which helps to detect uniquely matched reads for low-coverage genomes.

### 2.5 Generation of KMCP databases

Although KMCP is efficient and convenient in building custom databases, we provide three prebuilt databases (Supplementary Table S6). For bacteria and archaea, the representative GTDB r202 (Parks *et al*., 2022) species datasets including 47,894 genomes are used to build the KMCP database with the following parameters: sequence name filter = “plasmid”, *k* = 21, number of genome chunks = 10, chunk overlap = 150 bp, number of hash functions = 1, false-positive rate of Bloom filters = 0.3, and number of index files = 32. We choose the genome chunk number 10 for a balance of taxon identification accuracy and analysis time (Supplementary Fig. S6a and S6b) and chunk overlap of 150 bp for searching with common short reads of 150 bp in a single-end mode which has higher accuracy (Supplementary Fig. S6c). Genome_updater 0.2.5 (https://github.com/pirovc/genome_updater) is used to download fungal and viral genomes from the RefSeq or GenBank database. Four hundred and three fungal genomes from RefSeq r208 are used to construct the fungal database with the same parameters as the GTDB database. For GenBank r246, a maximum of five genomes are retained for each viral TaxId, leaving 27,936 viral genomes to build the viral database in which a lower false-positive rate of Bloom filters (0.05) is used in consideration of the small genome size. A snapshot of NCBI Taxonomy taxdump files on Dec 6, 2021, is used.

## 3 Results

### 3.1 Evaluation based on CAMI marine datasets

We evaluated KMCP against the ten simulated marine datasets from the second round of CAMI challenges compared with the top seven performant metagenomic profiling tools (Meyer *et al*.,2022), including mOTUs 2.5.1, MetaPhlAn 2.9.22, Centrifuge 1.0.4beta, DUDes 0.08, Metalign 0.6.2, Bracken 2.2, and CCMetagen 1.1.3 (Marcelino *et al*., 2020). The datasets were simulated from 155 newly sequenced marine isolate genomes and 622 public genomes from MarRef (Klemetsen *et al*., 2018). The numbers of species ranged from 256 to 381, and the abundances ranged from 0.001% to 11.533% (Supplementary Table S7 and Fig. S7a). KMCP built a database with 9,871 reference prokaryotic and 8,242 viral genomes, with a maximum of five genomes for each bacterial and archaeal species, from the same Jan 8, 2019, RefSeq snapshot used by other tools (Supplementary section S1.2). The gold standard profiles and results of different tools were downloaded and compared to those of KMCP using OPAL (Meyer *et al*., 2019).

KMCP showed taxon identification and relative abundance accuracies close to those of mOTUs2 and MetaPhlAn2 at the genus rank, while KMCP outperformed other tools at the species rank (Fig. 2a). The average completeness (recall, 0.915) of KMCP was slightly lower than that of Bracken (0.944) while higher than those of mOTUs2 (0.809), MetaPhlAn2 (0.850), and Centrifuge (0.887) at the species rank. For purity (precision), the value of KMCP (0.830) was lower than these of mOTUs2 (0.887) and DUDes (0.871). For relative abundance measured based on L1 norm error, KMCP had the second-lowest value at the genus rank and the lowest at the species rank. The weighted UniFrac error (Fig. 2b), which measures the similarity across taxonomic ranks between the gold standard and estimated abundances (Lozupone and Knight, 2005; Meyer *et al*., 2019), showed that KMCP had the lowest value. When we filtered out the predictions with the lowest relative abundances, totaling 0-10% within the species rank, the purity of all tools increased. However, the completeness decreased because the datasets were designed to have many low-abundance species (Fig. 2c, Supplementary Fig. S7a).

**Fig. 2.**
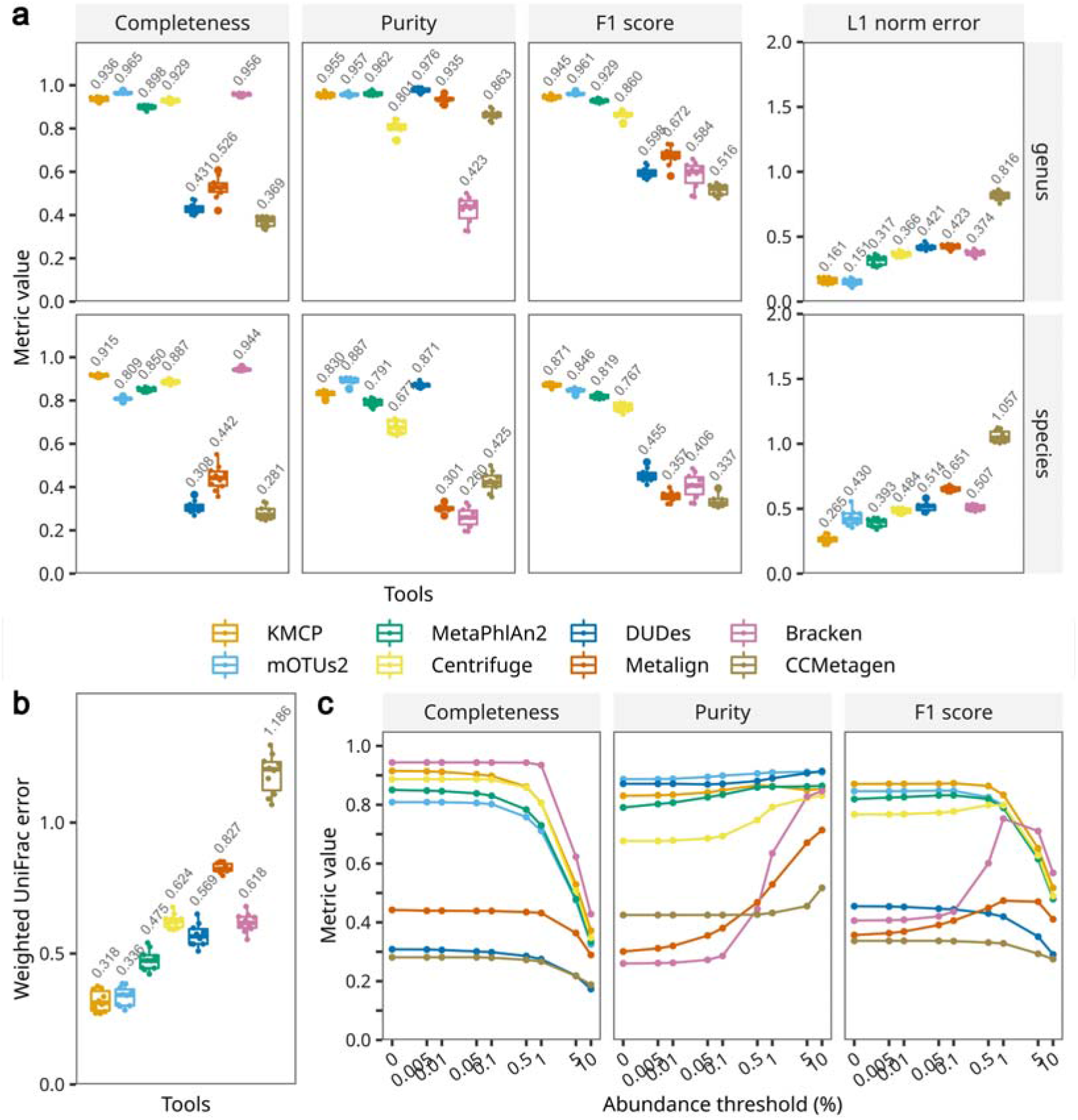
Taxonomic profiling accuracy based on CAMI marine datasets. **(a)**Assessment of taxon identification accuracy (completeness/recall, purity/precision, and F1 score) and relative abundance accuracy (L1 norm error) at the ranks of genus and species, with no abundance thresholding. **(b)** Relative abundance accuracy across all ranks, with no abundance thresholding. **(c)**Taxon identification accuracy changes at the species rank with abundance thresholds of 0 to 10%.

### 3.2 Evaluation based on simulated prokaryotic communities

We further evaluated KMCP alongside other taxonomy profilers of the newest versions with the latest databases, including Bracken 2.6.2, Centrifuge 1.0.4, Ganon 1.1.2, DUDes 0.08, SLIMM 0.3.4, MetaPhlAn 3.0.13 (Beghini *et al*., 2021), and mOTUs 3.0.1 (Ruscheweyh *et al*., 2022). Sun *et al*. (Sun *et al*., 2021) simulated twenty-five metagenomic sequence reads from five habitats with genomes selected from the intersection among the reference databases of MetaPhlAn2, mOTUs2, and Kraken2/Bracken (Supplementary section S1.3). Each dataset contained 3 Gb of 150-bp paired-end reads and presented a log-norm-distributed profile (Supplementary Fig. S7b), with numbers of species ranging from 21 to 113 and abundance ranging from 0.0277% to 48.0633% (Supplementary Table S7). To minimize database bias in the benchmarks, we built databases for Kraken, Bracken, Centrifuge, Ganon, DUDes, and SLIMM using the identical reference genomes in the KMCP database. Since MetaPhlAn3 and mOTUs3 are closely associated with official databases, the most recent databases (mpa_v30_CHOCOPhlAn_201901 for MetaPhlAn3 and v2.6.0 for mOTUs3) were used.

The *k*-mer-based methods (KMCP, Bracken, Centrifuge, and Ganon) and marker-gene-based methods (MetaPhlAn3 and mOTUs3) showed good completeness, with average values above 0.9 at both the genus and species ranks (Fig. 3a), while the alignment-based tools DUDes and SLIMM presented inadequate completeness. In terms of purity, KMCP, MetaPhlAn3, and mOTUs3 exhibited the highest values (above 0.9) at the genus rank, followed by Centrifuge (0.740). At the species rank, the average purity of mOTUs3 remained above 0.9, while those MetaPhlAn3 and KMCP dropped to 0.855 and 0.803, respectively. In terms of the F1 score, KMCP performed better than MetaPhlAn3 and Centrifuge at the genus rank, while the value is slightly lower than MetaPhlAn3 at the species rank. For L1 norm error, KMCP performed the best at the genus rank, followed by mOTUs and MetaPhlAn3; while at the species rank, mOTUs3 performed the best, followed by MetaPhlAn3 and KMCP. The Weighted UniFrac error results showed a similar trend to the L1 norm error at the genus rank (Fig. 3b). When low-abundance predictions were filtered out, the purities of Bracken, Centrifuge, and Ganon significantly increased but were still lower than those of mOTUs, MetaPhlAn3, and KMCP (Fig. 3c).

**Fig. 3.**
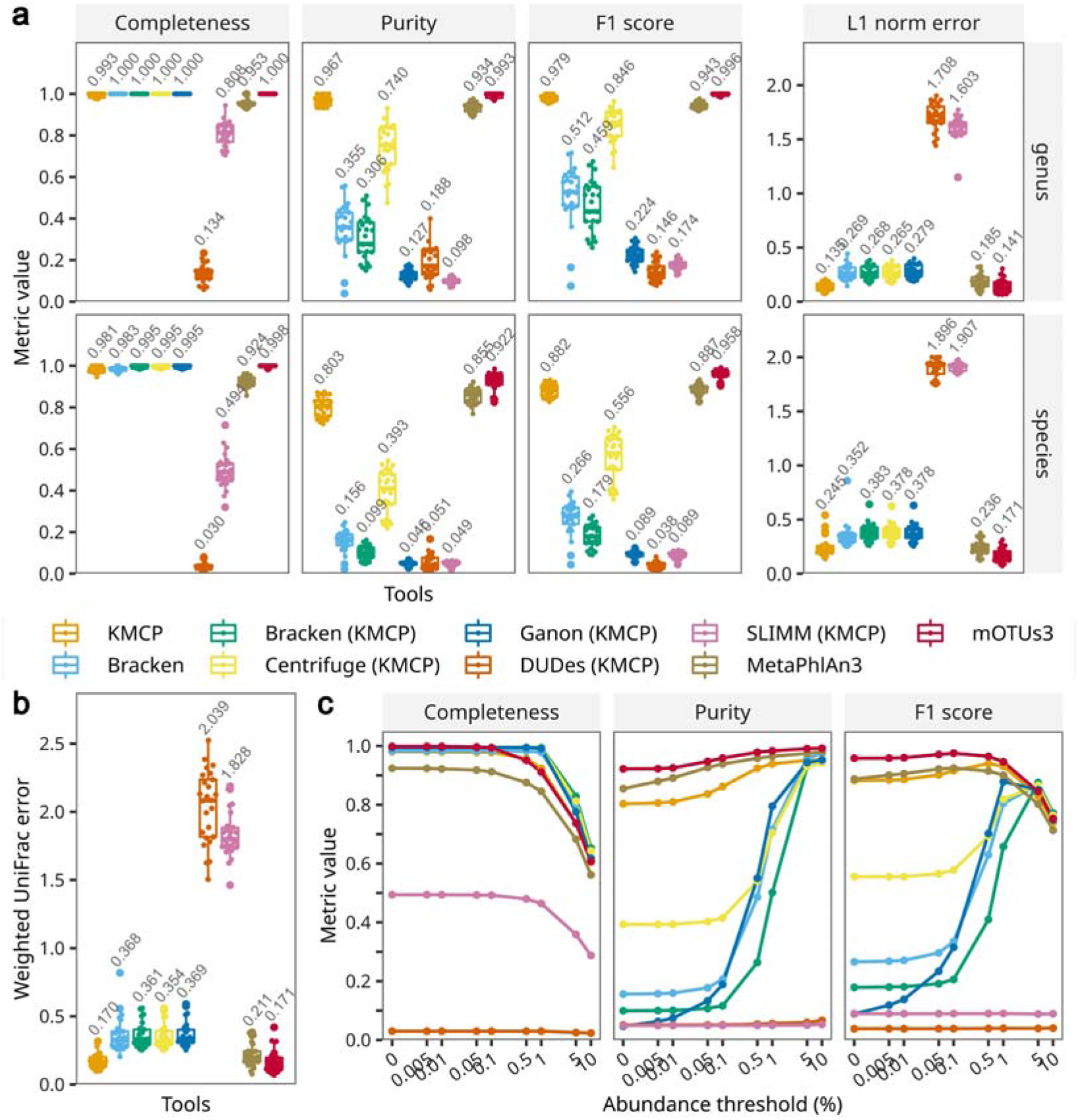
Taxonomic profiling accuracy based on simulated bacterial communities. Tools whose names are followed by (KMCP) use databases created with the same reference genomes used by KMCP. **(a)** Assessment of taxon identification accuracy (completeness/recall, purity/precision, and F1 score) and relative abundance accuracy (L1 norm error) at the ranks of genus and species, with no abundance thresholding. **(b)** Relative abundance accuracy across all ranks, with no abundance thresholding. **(c)** Taxon identification accuracy changes at the species rank with abundance thresholds of 0 to 10%.

### 3.3 Evaluation based on mock virome datasets

Roux *et al*. (Roux *et al*., 2016) designed two mock viral communities, including two ssDNA viruses and ten dsDNA viruses, with 16 datasets generated with three sequencing libraries. The numbers of species range from 10 to 12, and the abundances range from 0.002% to 74.9248% (Supplementary Table S7 and Fig. S7c). The clean data contained 50-250 bp of clean reads, and the total data ranged from 900 Mb to 2.5 Gb. mOTUs3, without virus detection support, was not included in this benchmark. MetaPhlAn3 was run with the option *“--add_viruses”* to add virus detection. KMCP, Bracken, and Centrifuge use databases constructed from identical reference genomes, including genomes of archaea, bacteria, fungi, and viruses (Supplementary section S1.4).

Since the mock viral communities were created by mixing viral capsids obtained from lysates, undigested bacterial DNA might be retained in the sequencing data. However, the relative abundances at the superkingdom rank showed variable results (Fig. 4a). KMCP and MetaPhlAn3 showed higher average viral abundances (99.72% and 99.02%), while the values generated by the other tools were below 80%. Then, we removed non-viral predictions and recomputed the taxonomic profiles using the *cami-filter* command in TaxonKit. KMCP, Bracken, Centrifuge, and Ganon, with the same reference genomes, showed completeness values equal to or higher than 0.9 at both the genus and species ranks (Fig. 4b). Bracken, with the official database, and MetaPhlAn3 presented much lower completeness, though they used databases built with reference genomes obtained after the publication of the sequencing data. For purity, the values of KMCP and Centrifuge were above 0.95 at the genus rank, while they decreased to 0.876 and 0.691, respectively, at the species rank, and the purity values of Bracken (KMCP) decreased from 0.857 to 0.369. Regarding taxon identification, KMCP ranked first at the species rank, with an average F1 score of 0.891, followed by Centrifuge, with a score of 0.814. KMCP also showed the best abundance estimation accuracy across taxonomic ranks with much lower L1 norm error and weighted UniFrac error (Fig. 4b and 4c).

**Fig. 4.**
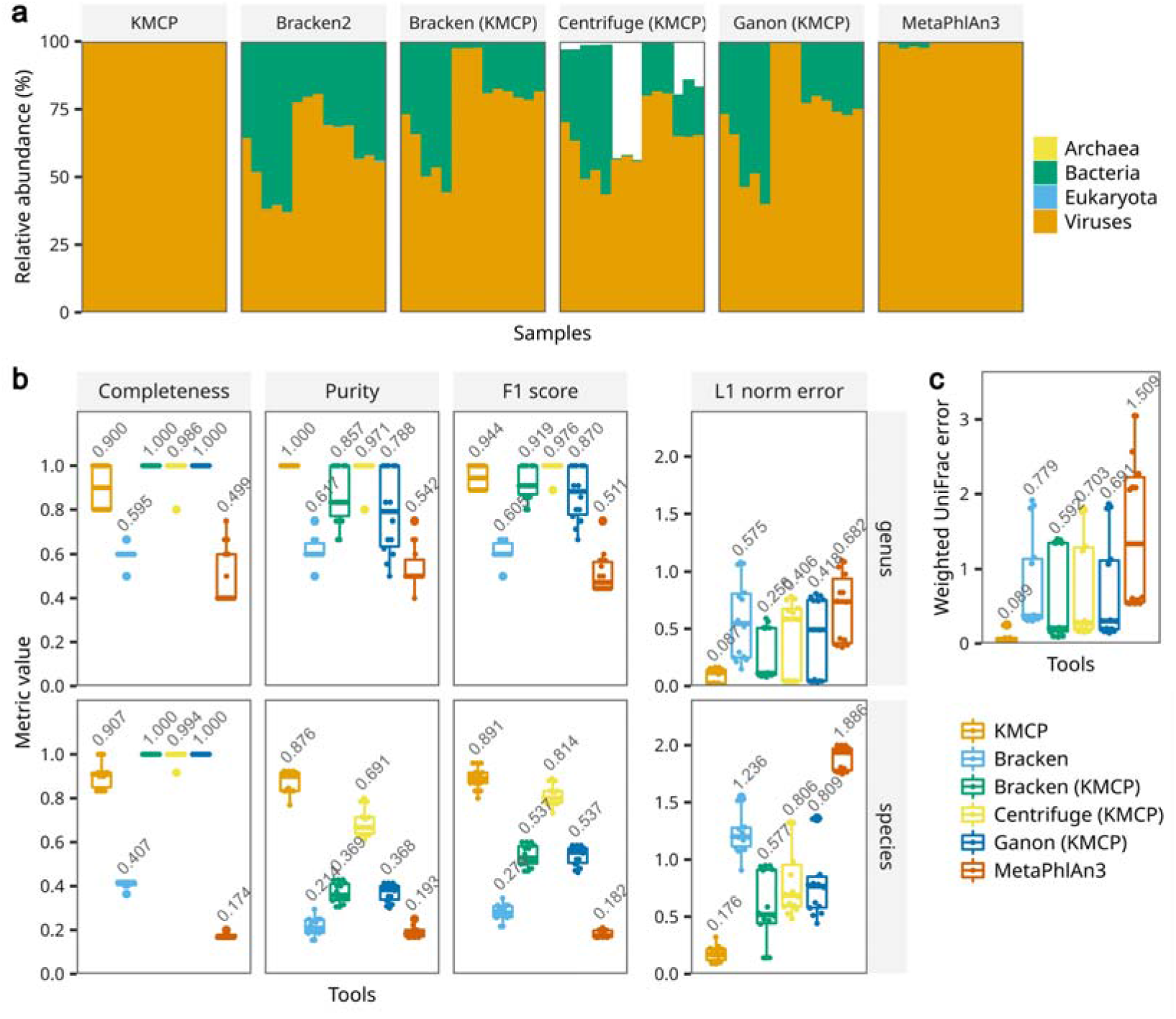
Taxonomic profiling accuracy based on mock virome communities. **(a)**Unfiltered microbial composition predicted by all the tools. **(b)**Assessment of viral presence-absence (completeness/recall, purity/precision, and F1 score) and relative abundance (L1 norm error) accuracy at the genus and species ranks, with no abundance thresholding. **(c)** Relative abundance accuracy of viruses across ranks, with no abundance thresholding. Tools whose names are followed by (KMCP) use databases constructed with the same reference genomes as KMCP.

We also compare KMCP against established viral detection and annotation software PhaGCN (Shang *et al*., 2021) and VIRify (Rangel-Pineros *et al*., 2022) (Supplementary section S1.4). The taxon identification accuracies at the family rank (the lowest common supported rank by the three tools) showed that all tools had a precision of 1, while KMCP had the highest average recall (0.971), followed by PhaGCN (0.600) and VIRify (0.543).

### 3.4 Evaluation of pathogen detection in clinical samples

The above results indicated that KMCP could accurately detect both prokaryotic and viral organisms in simulated and real metagenomic communities. We next evaluated whether KMCP could detect pathogens from low-depth clinical samples. Gu *et al*. (Gu *et al*., 2021) collected 182 body fluid samples from 160 patients as residual samples after routine clinical testing in a microbiology laboratory; among these samples, 87 with accessible short-read datasets that were verified by culture or 16S rRNA gene qPCR were analyzed with KMCP, Kraken2, Bracken, Centrifuge, and Ganon. Among these 87 samples, seventy-six were positive with one or more pathogens, and eleven were negative. The number of reads in the clean data ranged from 49 to 252,311 (Supplementary Table S12). MetaPhlAn3 and mOTUs3 were not tested in this case due to the limited ability of marker-gene-based methods to detect pathogens with ultralow coverage. According to our results, Bracken generally assigned more reads to the detected pathogens than Kracken2 alone, which did not affect the accuracy. Therefore, we ran Bracken immediately after Kraken2 on all samples. All tools were run according to the previous analysis but only required a minimum number of two reads for a prediction, and KMCP used the pathogen detection mode (Supplementary section S1.5).

The results (Table 1) showed that Bracken presented a significantly improved accuracy when using a more extensive database. Bracken, Centrifuge, and Ganon with the KMCP reference genomes all achieved a sensitivity of 80.26%, and KMCP, Bracken, and Ganon all presented a perfect specificity of 100%. The sensitivity of KMCP was slightly lower than those of Bracken, Centrifuge, and Ganon, with one misclassified sample (S19) (Supplementary Table S12). The low specificity of Centrifuge was due to the false-positive detection of a fungus (*Coccidioides immitis*) in three samples (S33, S35, S82).

**Table 1.** Accuracy of pathogen detection in clinical samples.

| Tools | Sensitivity | Specificity |
| --- | --- | --- |
| KMCP | 78.95% | 100% |
| Bracken | 73.68% | 81.82% |
| Bracken (KMCP) | 80.26% | 100.00% |
| Centrifuge (KMCP) | 80.26% | 72.73% |
| Ganon (KMCP) | 80.26% | 100.00% |

On the other hand, Bracken, Centrifuge, and Ganon mainly reported the detected organisms and associated read numbers. In contrast, KMCP reported more metrics than the read count, including the genome chunk fraction and similarity score (the 90^th^ percentile of *k*-mer coverage of all uniquely matched reads), which were also used to sort the predictions in the report (Supplementary section S1.5). These additional metrics allowed KMCP to provide confident results, with fewer predictions (**Fig. 5**a) and a higher mean rank of the pathogens in the prediction list (**Fig. 5**b). For predictions with small numbers of reads, the genome chunk fraction in KMCP results could also provide valuable evidence for determining the existence of the pathogens. For example, predictions with only 4-10 reads could still have genome chunk fractions ranging from 0.4 to 0.5 (**Fig. 5**c).

**Fig. 5.**
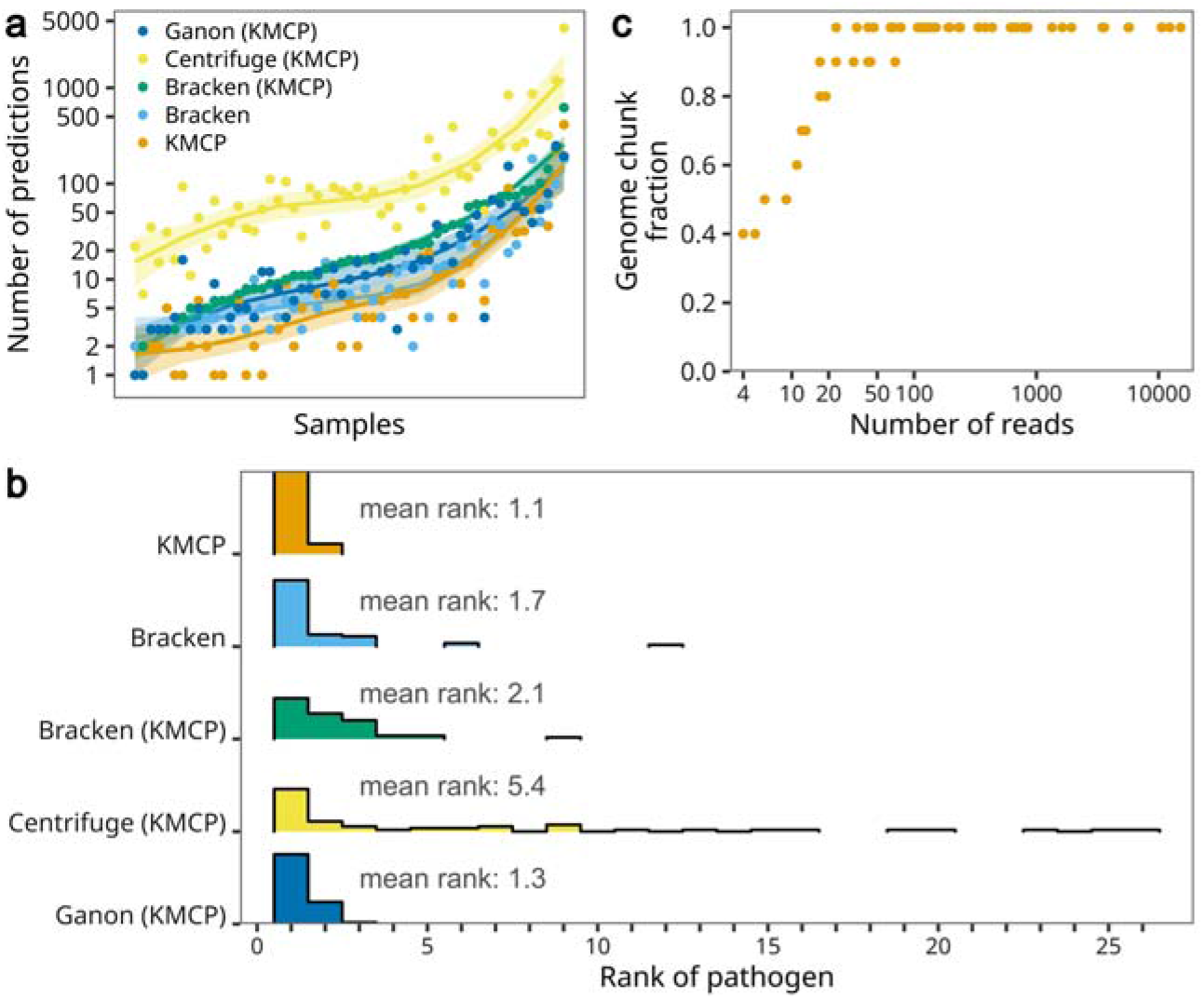
**(a)** The number of predictions in 55 samples where all tools correctly detected pathogens. Samples were sorted by the number of predictions with Bracken (KMCP). **(b)** Histogram of pathogen ranks in the 55 taxonomic profiling results. **(c)** The number of reads and the genome chunk fraction in the 60 true-positive predictions of KMCP. Tools whose names are followed by (KMCP) use databases constructed with the same reference genomes as KMCP.

### 3.5 Benchmarking computational requirements

We measured the runtime and maximum memory usage of database building and taxonomic profiling among KMCP and the above methods (specifications of the computing environments are available in Supplementary section S1.6). Since MetaPhlAn3 and mOTUs3 database construction requires many manual steps (Beghini *et al*., 2021; Milanese *et al*., 2019), we only created databases for KMCP, Kraken2, Bracken, Centrifuge, Ganon, DUDes, and SLIMM using 40 threads. We assessed the database building efficiency in terms of database size, building time, and peak memory occupation. The results illustrated that KMCP was much less time-consuming, required less memory than the other methods, and showed a smaller database size than Kraken2, Centrifuge, Ganon, DUDes, and SLIMM (Table 2 and Supplementary Table 11).

**Table 2.** Database size and building time and memory. The building time of KMCP includes the time of the *compute* and *index* commands.

| Tool | Index size (GB) | Building time | Peak memory (GB) |
| --- | --- | --- | --- |
| KMCP | 66 | 23 m | 15 |
| Bracken | 304 | 10 h 08 m | 258 |
| Centrifuge | 97 | 44 h 35 m | 522 |
| Ganon | 195 | 43 m | 197 |
| DUDes | 318 | 40 h 48 m | 922 |
| SLIMM | 317 | 40 h 34 m | 922 |
| MetaPhlAn3 | 3 | / | / |
| mOTUs3 | 8 | / | / |

Then, eight samples from the CAMI mouse gut metagenome datasets, each with 16.5 million paired-end 150-bp (5 Gb) reads, were used to assess metagenomic profiling time and memory requirements. All tools used 40 threads for taxonomic profiling (parameters of the tools are available in Supplementary section S1.6). The results (Fig. 6) showed that the marker-gene-based methods, including MetaPhlAn3 and mOTUs3, balanced analysis time and memory usage, while the other tools occupied much more memory. KMCP occupied a modest amount of memory (57.3 GB), but its speed was much slower than those of the other tools, with most of the time spent in reads searching. Bracken, Centrifuge, and Ganon, which used databases built with the same reference genomes as KMCP, achieved fast profiling but required more memory, with Bracken occupying more than 250 GB of memory. DUDes and SLIMM both used Bowtie2 for read mapping, which had an index of more than 300 GB (Table 2) and needed more than 200 GB of memory for mapping. The profiling step of DUDes was slower than that of SLIMM. Since KMCP has a scalable searching mode, we ran KMCP in a computer cluster with 24 computation nodes, referring to KMCP_HPC, where each node searched against a small database built with a partition of the reference genomes using 32 threads (the maximum number of available CPU threads in each node). The analysis time of KMCP decreased from 494.2 minutes to 28.7 minutes under these conditions.

**Fig. 6.**
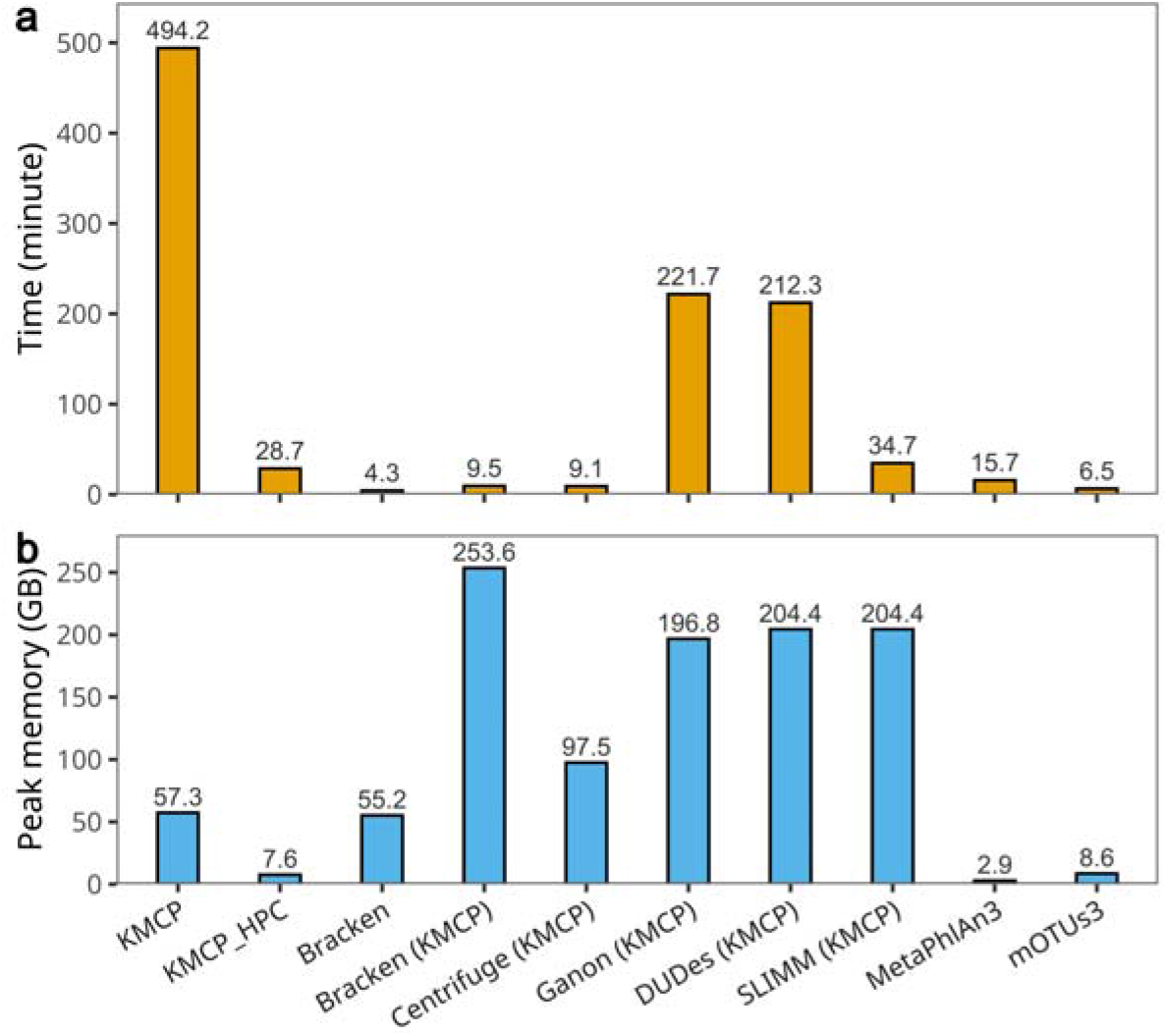
Per-sample runtime and maximum memory use for metagenomic profiling. The time of KMCP includes the time of the *search* and *profile* commands. KMCP_HPC refers to KMCP running on a computer cluster with each node performing searches against a small database built with a part of the reference genomes. Tools whose names are followed by (KMCP) use databases constructed with the same reference genomes as KMCP.

## 4 Discussion

The second round of CAMI challenges compared 22 methodological versions, and the results showed that the taxonomic profiling tools performed well among bacterial species. However, the accuracy was lower for archaea and viruses (Meyer *et al*., 2022), indicating a need for better reference sequence collection and tool development. Most microbiome studies have focused mainly on bacterial, archaeal, and fungal populations, with less attention paid to viral communities. However, bacteriophages play essential roles in microbiome ecology (Liang and Bushman, 2021; Townsend *et al*., 2021). Current viral metagenomic analysis approaches mainly rely on sequence assembly based on viral metagenomic sequencing data, while sample processing and genome assembly are challenging (Santiago-Rodriguez and Hollister, 2020). With the establishment of an increasing number of human gut prokaryotic (Almeida *et al*., 2021; Chibani *et al*., 2022; Hiseni *et al*., 2021) and virus/phage (Camarillo-Guerrero *et al*., 2021; Gregory *et al*., 2020; Nayfach *et al*., 2021) catalogs, it is possible to employ reference-based taxonomic profiling tools to investigate archaea, bacteria, and viruses simultaneously.

In this context, we have designed and implemented KMCP, a novel *k*-mer-based taxonomic profiling tool. *K*-mer-based methods apply fast exact *k*-mer matching and lack classification sensitivity for distant homologous genomes. Kraken2 uses the spaced seed (Brinda *et al*., 2015) to increase classification sensitivity. In KMCP, we use a small *k* value of 21 and retain all matches with *k*-mer coverage ≥ 0.55 by default, without LCA computation. A *k*-mer coverage of 0.55, approximately equal to a sequence identity of 96.5%, is chosen to balance the sensitivity of sequence searching and the generation of false positives resulting from the probabilistic data structure. Users can also apply a lower threshold by constructing a database with a lower false-positive rate at the expense of a larger index size. To minimize the false positives resulting from the lower *k*-mer coverage threshold, we used a three-round filtering step, with criteria including the *k*-mer similarity and genome chunks coverage information, to filter out suspicious matches. SLIMM also uses reference genome coverage information to remove unlikely genomes (Dadi *et al*., 2017) while it maps reads into bins of specific width (e.g., 1 kb) after the read mapping step. In contrast, KMCP splits reference genomes into predefined numbers (10 by default) of chunks in database building.

Benchmarking results based on simulated and real data indicate that KMCP accurately performs taxonomic profiling in both prokaryotic and viral populations and provides confident pathogen detection in infectious clinical samples. When applied to the simulated CAMI 2 marine metagenome datasets, KMCP presented the highest average F1 score at the species rank, and its L1 norm error value was the lowest. When applied to the prokaryotic datasets simulated with genomes selected from the intersection among the reference databases of MetaPhlAn2, mOTUs2, and Kraken2/Bracken reported in Sun *et al*., KMCP presented a slightly higher average F1 score than MetaPhlAn3 at the genus rank, and the average L1 norm error was the lowest. While at the species rank, KMCP had a slightly lower average F1 score than MetaPhlAn3. If using profiling mode 4, KMCP could increase the purity by 0.1 at the cost of a 0.04 decrease in completeness (Supplementary Table S9), but the F1 score would still be lower than that of mOTUs3. We note that prokaryotic benchmark datasets of CAMI2 marine and Sun *et al*. were simulated in distinct ways. The former used both public and newly sequenced genomes, while the latter used genomes from the intersection among the reference databases of MetaPhlAn2, mOTUs2, and Kraken2/Bracken; this might explain the inconsistent performance of the tools. Besides, the number of species and range of abundance varied, with the CAMI2 marine datasets having more (≥200) species and wider (0.001% to 11.533%) abundance ranges (Supplementary Table S7 and Fig. S7).

The benchmarking results based on the real mock virome datasets showed that KMCP outperformed Centrifuge, Bracken, and MetaPhlAn3, with higher taxon identification and abundance estimation accuracies. Though mOTUs3 showed the best accuracy based on prokaryotic metagenome datasets, it lacked the ability to detect viruses and was not included in this benchmark. For the other marker-gene-based method, MetaPhlAn3, we checked the database information and found that only 4 out of 12 viral species of the mock virome dataset were included, which partly explained the low average completeness (recall) of 0.174 at the species rank. KMCP also showed a better average recall at the family rank than two established viral detection and annotation methods, PhaGCN and VIRify. The lower recalls of these assembly-based strategies were probably due to the low coverage of some genomes (Supplementary Table S7 and Fig. S7c), which were insufficient for assembly.

We also showed that KMCP could detect low-coverage pathogens, including bacteria, fungi, and viruses, with as few as four reads in infectious clinical samples. KMCP presented a specificity of 100% and a sensitivity of 78.95%, slightly lower than the 80.26% sensitivity of Kraken2/Bracken and Ganon. In addition, KMCP generated shorter prediction lists and identified pathogens in priority positions by considering both sequence similarity and the genome chunk fraction, which could help researchers rapidly interpret the reports. Furthermore, the database size and memory occupation of KMCP were much lower than those of Kraken2/Bracken, Centrifuge, and Ganon with the same reference genomes.

Extensive and comprehensive reference databases of high quality are essential for producing good metagenomic results, and the increasing number of references imposes challenges regarding database size and construction efficiency among taxonomic profilers (Breitwieser *et al*., 2019). The searchable data structure BIGSI was invented to efficiently search for plasmid and resistance gene sequences in an index containing 447,833 microbial genomes (Bradley *et al*., 2019). Later, COBS improved BIGSI in terms of index size and searching speed. KMCP, reported in this paper, reimplements and modifies the COBS data structure. The index size of KMCP is smaller than that of COBS, and the batch searching speed is increased by ten times. However, after splitting the reference genomes into chunks, KMCP presents an *O(nK)* query time compared to the *O(K)* query time of BIGSI/COBS, where *K* is the number of reference genomes, and *n* is the number of genome chunks for each reference genome. We also tried implementing the RAMBO data structure (Gupta *et al*., 2019) (similar to BIGSI but closer to SBTs) in KMCP, which presents a sublinear query time at the cost of memory space and accuracy. However, the much larger index size did not improve the query efficiency in standalone machines, and the higher false-positive rate of queries negatively impacted the profiling accuracy. Fortunately, KMCP could utilize computer clusters to linearly accelerate read searching, with each node searching against a small database constructed with a partition of the reference genomes.

Ganon and KMCP are both efficient in indexing a large number of reference genomes. IBF, a data structure very similar to BIGSI, is used in Ganon for indexing *k*-mers. Technically, IBF, BIGSI, and COBS all interleave each row of bits of Bloom filters and save them as a byte list in either database (BIGSI) or a binary file (IBF and COBS). Ganon uses a smaller *k*-mer size (19) than KMCP (21) by default, theoretically generating fewer *k*-mers. Ganon sets a low false-positive rate of 0.05 and uses three hash functions for Bloom filters to balance database size and classification accuracy. In KMCP, we use a high false-positive rate of 0.3 for creating smaller databases and use only one hash function to increase search speed. However, the benchmarking results based on both prokaryotic and viral datasets demonstrated that KMCP presents higher accuracies than Ganon in taxon identification, as the latter method shows low precision among low-abundance taxa.

The accuracy of taxonomic profiling relies on the reference genome and microbial taxonomy. The NCBI taxonomy database is a curated classification and nomenclature database that incorporates all organisms available in major public sequence databases, including GenBank and RefSeq (Schoch *et al*., 2020). However, with the continuous updating of the taxonomy database, TaxIds are added, deleted, and merged into other TaxIds; the taxonomic ranks, names, and lineage are also changed (Shen and Ren, 2021). The modular design of KMCP separates taxonomic information from the *k*-mer index, allowing the database to adapt to new taxonomy versions. As reference genomes, we chose representative species genomes from GTDB, which provides a standardized bacterial and archaeal taxonomy based on genome phylogeny, to build the prokaryotic database. Fungal genomes from RefSeq and viral genomes from GenBank were used to create fungal and viral databases, respectively. Based on the same reference genomes, KMCP generates a much smaller database and requires less memory for searching than Kraken2/Bracken, Centrifuge, and Ganon. Moreover, the lack of genome preprocessing (clustering or low-complexity-region masking) and the fast indexing speed of KMCP enable users to build custom databases efficiently, and scalable searching allows users to perform searches against multiple databases. To update databases, users can either rebuild the database with newly added reference genomes or just build an additional database for the new genomes because search results against multiple databases can be rapidly merged.

In summary, many factors affect the accuracy of taxonomic profiling, including the number of reference genomes, the diversity of the metagenomic communities, and the methodology of the software. The high efficiency in database building and the scalability in reads searching make KMCP efficiently utilize the increasing number of reference genomes. And KMCP has combined *k*-mer similarity and genome coverage information to improve taxonomic profiling accuracy in both prokaryotic and viral populations. There is still room for improvement. For prokaryotic metagenome datasets, KMCP showed a lower purity (precision) at the species rank than at the genus rank, and the abundance estimation at the species rank was less accurate than those of the marker-gene-based method, MetaPhlAn3 and mOTUs3. We consider it worthwhile to improve taxon identification and abundance estimation in future work.

## Acknowledgments

We thank Zhi-Luo Deng (Department of Computational Biology, Helmholtz Centre for Infection Research, Braunschweig, Germany) for valuable advice on metagenomic profiling and benchmarking. We thank Robert Clausecker (Zuse Institute Berlin, Germany) for developing the high-performance vectorized positional popcount package (pospop) during the development of KMCP. We thank Jishan Tan (Department of Clinical Laboratory, General Hospital of Western Theater Command, Chengdu, China) for clinical microbiology diagnosis consultations. We thank Lei Zhang (Deepbiome Co. Ltd, China) for the valuable discussion on Virus identification and annotation. We thank Chenhao Li (Massachusetts General Hospital, Broad Institute, USA), Zhi-Luo Deng, Qi Zhao (Sun Yat-sen University Cancer Center, Guangzhou, China), and Yujie Zhao for advising and commenting on the manuscript.

