## Supplementary materials for "KMCP: accurate metagenomic profiling of both prokaryotic and viral populations by pseudo-mapping"

### Index of Tables

Figure S8. Taxon identification accuracies at the family rank on the mock virome dataset ... 20

### 1 Methods

#### 1.1 Optimization of searching speed in the modified COBS index

Many optimizations are performed to accelerate batch querying in the modified COBS index. Multiple hashing functions are optionally used to reduce the false-positive rate of Bloom filters, which requires a bitwise AND step to double-check the existence of a *k*-mer. Instead of the byte-by-byte operation performed in COBS, the AND step is optimized with Single-Instruction Multiple-Data (SIMD) instructions. For the ADD step, which cumulatively adds the number of matched *k*-mers in a Bloom filter, COBS uses array lookup to avoid unpacking the bit-packed count information. However, array lookup requires frequent memory access, which is fast but not free. Here, we use a high-performance vectorized positional *popcount* method (https://github.com/clausecker/pospop), which utilizes SIMD instructions to unpack and count *k*-mers in CPU registers for a list of bytes. To balance the speed of memory access for collecting column-wise bytes and invoking the *pospop* function, we set the size of the byte list to 64 (i.e., counting matched elements for 64 *k*-mers in 8 (8 bits for a byte) genome chunks in one function invocation). In addition, sixty-four is the cache line size for most 64-bit computers, which helps to reduce cache misses and provides extra speedup. For the left *k*-mers (<64) of the query sequence, we unroll the loops of 63 possible *k*-mer sizes to increase CPU instruction execution efficiency. Furthermore, many language-level optimizations are performed to reduce memory occupation and accelerate searching speed. A complete list of differences in algorithm and data structure between KMCP and COBS is listed in Table S4.

#### 1.2 Benchmarking KMCP using the CAMI framework

Snapshots of RefSeq and NCBI Taxonomy dump files on Jan 08, 2019, are used to create the KMCP database. TaxIds of archaea and bacteria are retrieved with TaxonKit 0.10.1 (Shen and Ren, 2021) and used to extract genomes from the RefSeq snapshot. The assembly accessions of all genomes are sorted in alphabetical order, and a maximum of five genomes of a species are retained. Since the RefSeq snapshot only contains prokaryotic genomes, we downloaded the viral assembly summary file from RefSeq FTP on Nov 11, 2021, and extracted viral references before Jan 08, 2019. Finally, 9,871 prokaryotic and 8,242 viral genomes are used to build KMCP databases with the parameters described in the "Generation of the KMCP databases" section. The 10 marine datasets were download from https://data.cami-challenge.org/participate. KMCP v0.9.0 uses the command "*kmcp search -j 40 -d $db read.fq.gz -o read.kmcp@$db.tsv.gz*" for searching reads against prokaryotic and viral databases, command of "*kmcp merge read.kmcp@*.tsv.gz -o read.kmcp.tsv.gz*" for merging search results, and command of "*kmcp profile -m 3 -X $taxdump -T $taxidmap read.kmcp.tsv.gz -o read.kmcp.tsv.gz.k-m3.profile -C read.kmcp.tsv.gz.c-m3.profile --show-rank superkingdom,phylum,class,order,family,genus,species -s $sampleID*" for profiling. The gold standards and profiles of other tools are download from https://zenodo.org/record/5006866, and OPAL (commit edb6f41 in dev branch) (Meyer *et al.*, 2019) is used to assess taxonomic metagenome profilers. R 4.1.2 (R Core Team, 2022), ggplot2 3.3.5 (Wickham, 2016), ggbreak 0.0.9 (Xu *et al.*, 2021), and some other R packages are used for data visualization.

#### 1.3 Benchmarking KMCP using simulated bacterial communities

Sun *et al*. (Sun *et al.*, 2021) simulated twenty-five simulated metagenomic sequence reads from five habitats with genomes selected from the intersection among the reference databases of MetaPhlAn2, mOTUs2, and Kraken2. Each dataset contains 3 Gb of 150-bp paired-end reads. KMCP 0.9.0, with the standard database (Dec 6, 2021); Kraken 2.1.2 and Bracken 2.6.2, with the PlusPF collection of the official databases (May 17, 2021); MetaPhlAn 3.0.13, with the mpa_v30_CHOCOPhlAn_201901 database (July 22, 2020); and mOTUs 3.0.1, with the 2.6.0 database (June 16, 2021), are used for benchmarking. In addition, the genomes for building the KMCP database, including 47,894 prokaryotic genomes from GTDB, 27,936 viral genomes from GenBank, and 403 fungal genomes from RefSeq, are used to create databases for Kraken2/Bracken, Bracken, Centrifuge 1.0.4, Ganon 1.1.2, DUDes 0.08, and SLIMM 0.3.4 with the default parameters. SeqKit 2.1.0 (Shen *et al.*, 2016) is used in reference genome preparation.

KMCP uses the command "*kmcp search -j 40 -d $db read1.fq.gz read2.fq.gz -o read.kmcp@$db.tsv.gz*" for searching reads against prokaryotic and fungal databases, command of "*kmcp merge read.kmcp@*.tsv.gz -o read.kmcp.tsv.gz*" for merging search results, and command of "*kmcp profile -m 3 -X $taxdump -T $taxidmap read.kmcp.tsv.gz -o read.kmcp.tsv.gz.k-m3.profile -C read.kmcp.tsv.gz.c-m3.profile --show-rank superkingdom,phylum,class,order,family,genus,species -s $sampleID*" for profiling. MetaPhlAn3 uses the command "*metaphlan --nproc 40 -x $db --bowtie2db $dbdir --input_type fastq read1.fq.gz,read2.right.fq.gz --bowtie2out bowtie2.bz2 --CAMI_format_output -o mpa3.profile*", and mOTUs3 uses the command "*motus profile -t 40 -f read1.fq.gz -r read2.fq.gz -p -C precision -o motus.profile*". Kraken2 uses the command "*kraken2 --threads 40 --db $db --memory-mapping --gzip-compressed --paired read1.fq.gz read2.fq.gz --report read.kreport > /dev/null*" for classification, and Bracken uses the command "*for r in S G F O C P D; do est_abundance.py -k $db/database150mers.kmer_distrib -l $r -t 10 -i read.kreport -o read.bracken.level-$r; done*" to estimate abundance at species, genus, family, order, class, phylum, and superkingdom ranks, the script tocami.py from CAMI (https://github.com/hzi-bifo/cami2_pipelines/blob/master/bin/tocami.py, commit a0e2ec2) is used to convert the concatenated abundance tables to the CAMI taxonomic profiling format. Centrifuge uses the command "*centrifuge -t -p 40 --mm -x $db -q -1 read1.fq.gz -2 read2.fq.gz --report-file read.cf-report.tsv -S read.cf.tsv*" for classification and uses the command "*centrifuge-kreport -x $db read.cf.tsv > read.kreport*" to generate Kraken-style report, which is further converted to CAMI format with the tocami.py script. Ganon uses the command "*ganon classify -t 40 -d $db -p read1.fq.gz read2.fq.gz -o read*" for classification and the command "*ganon table -l percentage --header taxid -r species -I read.tre -o read.tsv*" to generate the composition table. DUDes and SLIMM use the same command, "*bowtie2 --mm -p 40 -x $bowtie2db --no-unal --very-fast -k 100 -q -1 read1.fq.gz -2 read2.fq.gz -S read.sam*", for read mapping. DUDes uses the command "*DUDes.py -t 1 -s read.sam -d $db -o read*", and SLIMM uses the command "*slimm -w 1000 -o read $db read.sam*" for profiling. The taxonomic profiles of the gold standards and all taxonomic profiles were converted to CAMI format with the NCBI taxonomy dump file of Dec 6, 2021, and the virus predictions were filtered using TaxonKit 0.10.1.

#### 1.4 Benchmarking KMCP using mock virome communities

Roux *et al*. (Roux *et al.*, 2016) designed two mock viral communities including two ssDNA viruses and ten dsDNA viruses. Each virus was grown on its specific host (*Pseudoalteromonas*, *Cellulophaga baltica,* or *Escherichia coli*), and viral capsids were obtained from lysates and mixed. Sixteen datasets were generated with three sequencing libraries. The clean data contained paired and unpaired reads with lengths of 50 to 250 bp, and the amount of data from each sample ranged from 900 Mb to 2.5 Gb. The normalized coverage of each genome was used as a proxy for the relative abundance of each viral genome (Roux *et al.*, 2016). mOTUs3, without virus detection support, was not included in this benchmark. MetaPhlAn3 was run with the flag "*--add_viruses*" to add virus detection. KMCP v0.9.0 searches against prokaryotic, fungal, and viral databases. After the initial community composition analysis, nonviral predictions were filtered out using TaxonKit 0.10.1.

Reads are assembled with SPAdes 3.15.5 (Nurk *et al.*, 2017) with the metagenomic modes (--meta) and a *k*-mer size of 55, and contigs shorter than 2 kb are filtered out. VIRify 0.4.0 (Rangel-Pineros *et al.*, 2022) is used to perform virus detection and annotation from the contigs with the command of "*nextflow run EBI-Metagenomics/emg-viral-pipeline -r v0.4.0 --databases $db --cachedir $cache --ncbi $ete3tax --fasta $contigs --workdir work --output virify --length 2 -profile local,singularity*" with the same taxonomy data (NCBI taxonomy dump file of Dec 6, 2021,) to KMCP. DeepVirFinder v1.0 (Ren *et al.*, 2020) is used to identify viral sequences with the default parameters and then PhaGCN (commit 5c40b59) (Shang *et al.*, 2021) is used to annotate viral sequences with the default parameters. At last, the accuracies of taxon identification at the family rank are computed.

#### 1.5 Benchmarking KMCP using infectious clinical samples

Gu *et al.* (Gu *et al.*, 2021) collected 182 body fluid samples from 160 patients as residual samples after routine clinical testing in the microbiology laboratory, among which 87 have accessible short-read datasets and have been verified by culture or 16S rRNA gene qPCR. The number of reads in the clean data ranges from 49 to 252,311 (Table S3). Kraken2 and Bracken, Centrifuge, and KMCP are used to detect pathogens, and all tools require a minimum number of two reads. KMCP uses a preset profiling mode with the highest sensitivity for pathogen detection, in which the predictions are sorted in reverse order by the product of the genome chunk fraction and the similarity score, rather than the read number or taxonomic abundance, as in other tools. The score is the 90$^{th}$ percentile of *k*-mer coverage of all uniquely matched reads. For Bracken, the number of new estimated reads (column *new_est_reads*) is used for sorting. The number of reads (column *numReads*) is used in Centrifuge. Similarly, the taxon abundances in the Ganon results are sorted by the number of reads.

#### 1.6 Benchmarking runtime and storage requirement of database building and profiling

The performance of KMCP and other tools is analyzed on a server with four Intel(R) Xeon(R) Gold 6248 processors (80-CPUs/160-threads) and one terabyte of main memory, and the reference genomes and databases are stored on solid-state disks, while the sequencing reads are saved on hard disk drives. The operation system is Centos 7.9.2009 with the kernel of 3.10.0-1160.11.1.

The genomes used for building the KMCP databases, including 47,894 prokaryotic genomes from GTDB, 27,936 viral genomes from GenBank, and 403 fungal genomes from RefSeq, are used to create databases for Kraken, Bracken, and Centrifuge with the default parameters. MetaPhlAn3 and mOTUs3 are tightly associated with the official databases, and the most recent databases (mpa_v30_CHOCOPhlAn_201901 for MetaPhlAn3 and 3.0.1 for mOTUs3) are used. Eight samples from the CAMI mouse gut metagenome datasets are used to assess the analysis time and memory requirement. All tools use 40 threads for database building and taxonomic profiling. The scaling test of KMCP is performed on a computer cluster with 24 nodes, each with one Hygon C86 7185 32-core processor and 124 GB of memory. A script (https://github.com/shenwei356/easy_sbatch) is used for the batch submission of search jobs against all databases.

### 2 Availability of data and materials

The CAMI2 marine datasets, mouse gut datasets, and the Jan 8, 2019, RefSeq snapshot can be downloaded from https://data.cami-challenge.org/participate, and the gold standard taxonomic profile and profiles of other tools can be downloaded from https://zenodo.org/search?page=1&size=20&q=keywords:%22CAMI%202%20Mouse%20Gut%20Toy%20data%20set%22&keywords=taxonomic%20profiling. The reference genomes and database used by KMCP can be downloaded from https://1drv.ms/u/s!Ag89cZ8NYcqtjVVADr8r--fnKFt-?e=ivNZNK. Simulated metagenomic sequence reads from Sun *et al.* (Sun *et al.*, 2021) can be downloaded from https://figshare.com/projects/Pitfalls_and_Opportunities_in_Benchmarking_Metagenomic_Classifiers/79916, and the gold standard profiles in CAMI format can be downloaded from https://github.com/shenwei356/sun2021-cami-profiles/releases/tag/v2021-12-06. Mock virome communities A and B from Roux *et al.* (Roux *et al.*, 2016) can be downloaded from https://datacommons.cyverse.org/browse/iplant/home/shared/iVirus/DNA_Viromes_library_comparison, and the gold standard profiles in CAMI format can be downloaded from https://github.com/shenwei356/roux2016-mock-virome-cami-profile/releases/tag/v2021-12-06. The metagenomic samples of infected body fluids from Gu *et al*. (Gu *et al.*, 2021) can be downloaded from https://www.ncbi.nlm.nih.gov/bioproject/?term=PRJNA558701, and the list of run accessions can be downloaded from https://github.com/shenwei356/kmcp/blob/cb98c64c7dda4f1863c603a6dd38296572479396/benchmarks/real-pathogen-gu2020/acc2sample.tsv.

The source code of KMCP is under the MIT license and is publicly available on Github (https://github.com/shenwei356/kmcp) and Zenodo (https://zenodo.org/record/7117838). The prebuilt databases can be downloaded from https://1drv.ms/u/s!Ag89cZ8NYcqtjHwpe0ND3SUEhyrp?e=QDRbEC. Benchmarking steps, generated taxonomic profiles, plotting scripts, and other data generated and analyzed during the current study are available in the Zenodo repository (https://doi.org/10.5281/zenodo.7144333 ).

### 3 Tables

**Table S1. List of KMCP sub-commands.**

| Sub-command | Description |
|---|---|
| `compute` | Generating *k*-mers (sketches) from FASTA/Q sequences |
| `index` | Constructing database from *k*-mer files |
| `search` | Searching sequences against a database |
| `merge` | Merging search results from multiple databases |
| `profile` | Generating taxonomic profile from search results |
| `utils split-genomes` | Splitting genomes into chunks |
| `utils unik-info` | Printing information of .unik file |
| `utils index-info` | Printing information of index file |
| `utils ref-info` | Printing information of reference chunks in a database |
| `utils cov2simi` | Convert *k*-mer coverage to sequence similarity |
| `utils query-fpr` | Compute the maximal false positive rate of a query |
| `utils filter` | Filtering search results and finding species/assembly-specific queries |
| `utils merge-regions` | Merging species/assembly-specific regions |

**Table S2. The three options to create extra smaller block.** Uneven genome size distribution would make bloom filters of the last block with the most *k*-mers extremely huge, so three thresholds are used to split the last block.

| Option | Default threshold | Size of block | Description |
|---|---|---|---|
| -x/--block-sizeX-kmers-t | 10 M | 256 (settable) | If the number of *k*-mers of an .unik file exceeds the threshold, the block size is changed to 256. |
| -8/--block-size8-kmers-t | 20 M | 8 | If the number of *k*-mers of an .unik file exceeds the threshold, the block size is changed to 8. |
| -1/--block-size1-kmers-t | 200 M | 1 | If the number of *k*-mers of an .unik file exceeds the threshold, an individual index is created for this file. |

**Table S3. Comparison of database size and building time on GTDB dataset**. The 47,894 bacteria and archaea genomes from GTDB r202 representative collection were used to build databases. COBS and KMCP used the same parameters of k = 31, block size = 1024, hash functions = 1, false-positive rate of bloom filters = 0.3.

| Tool | Database size | Building time | Temporary files size |
|---|---|---|---|
| COBS | 86.96 GB | 29m 55s | **160.76 GB** |
| KMCP | **55.15 GB** | **21m 04s** | 935.11 GB |

**Table S4. The differences between KMCP and COBS.**

| Category | Iterm | COBS | KMCP | Comment |
|---|---|---|---|---|
| Algorithm | K-mer hashing | xxhash | ntHash | The xxHash is a general-purpose hashing function while the ntHash is a recursive hash function for DNA/RNA |
| | Bloom filter hashing | xxhash | Using *k*-mer hash values | Avoid hash computation |
| | Multiple-hash functions | xxhash with different seeds | Generating multiple values from a single one | Avoid hash computation |
| | Single-hash function | Same to multiple-hash functions | Separated workflow | Reducing loops |
| | AND step | Serial bitwise AND | Vectorized bitwise AND | Bitwise AND for >1 hash functions |
| | PLUS step | Serial bit-unpacking | Vectorized positional popcount with pospop | Counting from bit-packed data |
| Index structure | Size of blocks | / | Using extra thresholds to split the last block with the most *k*-mers | Uneven genome size distribution would make bloom filters of the last block extremely huge |
| | Index files | Concatenated | Independent | Index files |
| | Index loading | mmap, loading complete index into RAM | mmap, loading complete index into RAM, seek | Index loading |
| Input/ output | Input files | FASTA/Q, McCortex, text | FASTA/Q | Input files |
| | Output | Target and matched *k*-mers | Target, matched *k*-mers, query FPR, etc. | Output |

**Table S5. Main reference filtering criteria of the six profiling modes.**

| Filtering criteria | m0 | m1 | m2 | m3 (default) | m4 | m5 |
|---|---|---|---|---|---|---|
| Minimum reads of a chunk | 1 | 5 | 10 | 50 | 100 | 100 |
| Minimum chunk fraction | 0.2 | 0.6 | 0.7 | 0.8 | 1 | 1 |
| Maximum standard deviation of the relative depth of genome chunks | 10 | 2 | 2 | 2 | 2 | 1.5 |
| Minimum uniquely matched reads | 1 | 2 | 5 | 20 | 50 | 50 |
| Minimum high-confidence uniquely matched reads | 1 | 1 | 2 | 5 | 10 | 10 |
| Minimum *k*-mer coverage of high-confidence uniquely matched reads | 0.7 | 0.7 | 0.7 | 0.75 | 0.8 | 0.8 |
| Minimum proportion of high-confidence uniquely matched reads | 0.01 | 0.1 | 0.2 | 0.1 | 0.1 | 0.15 |
| Keeping main matches | Yes | | | | | |
| Maximum query coverage gap | 0.4 | | | | | |

**Table S6. Summary of KMCP databases.**

| Database | Source | #species | #assemblies | *k* | Chunks | FPR | Size (GB) |
|---|---|---|---|---|---|---|---|
| Bacteria and Archaea | GTDB r202 | 28073 | 47894 | 21 | 10 | 0.3 | 58.03 |
| Fungi | Refseq r208 | 398 | 403 | 21 | 10 | 0.3 | 4.18 |
| Viruses | GenBank r246 | 23632 | 27936 | 21 | 10 | 0.3 | 4.72 |

**Table S7. Summary of species in benchmark datasets, including the number of species and the percentage of abundance.**

| Dataset | Environment | Minimum #species | Maximum #species | Minimum %abundance | Maximum %abundance |
|---|---|---|---|---|---|
| CAMI marine datasets | / | 256 | 381 | 0.0010 | 11.5330 |
| Simulated prokaryotic communities | Building | 90 | 113 | 0.0370 | 15.7443 |
| | Gut | 56 | 76 | 0.0277 | 19.2673 |
| | Oral | 73 | 98 | 0.0364 | 12.2192 |
| | Skin | 52 | 80 | 0.0504 | 14.9236 |
| | VG | 21 | 38 | 0.1067 | 48.0633 |
| Mock virome datasets | / | 10 | 12 | 0.0020 | 74.9248 |

**Table S8. Accuracies of different profiling modes of KMCP on CAMI2 marine datasets**.

| Rank | Filtering criteria | m1 | m2 | m3 | m4 | m5 |
|---|---|---|---|---|---|---|
| Genus | Completeness | **0.9449** | 0.9388 | 0.9358 | 0.9274 | 0.9219 |
| | Purity | 0.9367 | 0.9528 | 0.9551 | 0.9562 | **0.9559** |
| | F1 score | 0.9408 | **0.9458** | 0.9454 | 0.9415 | 0.9385 |
| | L1 norm error | **0.1585** | 0.1626 | 0.1611 | 0.1828 | 0.2153 |
| Species | Completeness | **0.9298** | 0.9200 | 0.9151 | 0.9037 | 0.8955 |
| | Purity | 0.7488 | 0.8202 | 0.8303 | 0.8739 | **0.8976** |
| | F1 score | 0.8294 | 0.8672 | 0.8705 | 0.8885 | **0.8964** |
| | L1 norm error | 0.2639 | 0.2650 | 0.2653 | 0.2649 | **0.2665** |
| / | Weighted UniFrac error | 0.3191 | 0.3217 | **0.3183** | 0.3346 | 0.3573 |

**Table S9. Accuracies of different profiling modes of KMCP on simulated bacterial communities**.

| Rank | Filtering criteria | m1 | m2 | m3 | m4 | m5 |
|---|---|---|---|---|---|---|
| Genus | Completeness | **1.0000** | 0.9967 | 0.9926 | 0.9748 | 0.9687 |
| | Purity | 0.8238 | 0.9168 | 0.9667 | 0.9857 | 0**.9875** |
| | F1 score | 0.9021 | 0.9547 | 0.9794 | **0.9802** | 0.9779 |
| | L1 norm error | **0.1287** | 0.1356 | 0.1345 | 0.1604 | 0.1927 |
| Species | Completeness | **0.9904** | 0.9765 | 0.9806 | 0.9440 | 0.9062 |
| | Purity | 0.4851 | 0.6656 | 0.8033 | 0.9018 | **0.9282** |
| | F1 score | 0.6500 | 0.7902 | 0.8823 | **0.9221** | 0.9169 |
| | L1 norm error | **0.2417** | 0.2492 | 0.2453 | 0.2728 | 0.3090 |
| / | Weighted UniFrac error | **0.1617** | 0.1710 | 0.1696 | 0.2035 | 0.2509 |

**Table S10. Accuracies of different profiling modes of KMCP on mock virome communities**.

| Rank | Filtering criteria | m1 | m2 | m3 | m4 | m5 |
|---|---|---|---|---|---|---|
| Genus | Completeness | **0.9143** | **0.9143** | 0.9000 | 0.8821 | 0.8821 |
| | Purity | 0.9536 | 0.9714 | **1.0000** | **1.0000** | **1.0000** |
| | F1 score | 0.9263 | 0.9365 | **0.9444** | 0.9342 | 0.9342 |
| | L1 norm error | 0.0889 | 0.0889 | **0.0875** | 0.1532 | 0.1532 |
| Species | Completeness | **0.9311** | **0.9311** | 0.9073 | 0.8769 | 0.8769 |
| | Purity | 0.8441 | 0.8669 | 0.8757 | **0.9044** | **0.9044** |
| | F1 score | 0.8839 | **0.8966** | 0.8906 | 0.8898 | 0.8898 |
| | L1 norm error | 0.1501 | **0.1480** | 0.1762 | 0.2732 | 0.2732 |
| / | Weighted UniFrac error | 0.0848 | **0.0846** | 0.0885 | 0.1148 | 0.1148 |

**Table S11. Database size, building time and memory of KMCP for performance benchmark.**

| Database | Command | Index size | Building time | Peak memory |
|---|---|---|---|---|
| Bacteria and Archaea | compute | / | 9m 01s | 4.81 GB |
| | index | 56.94 GB | 10m 47s | 11.02 GB |
| Fungi | compute | / | 59.405s | 14.44 GB |
| | index | 4.16 GB | 44.349s | 1.36 GB |
| Viruses | compute | / | 22.920s | 4.54 GB |
| | index | 4.33 GB | 28.894s | 2.87 GB |

### 4 Figures

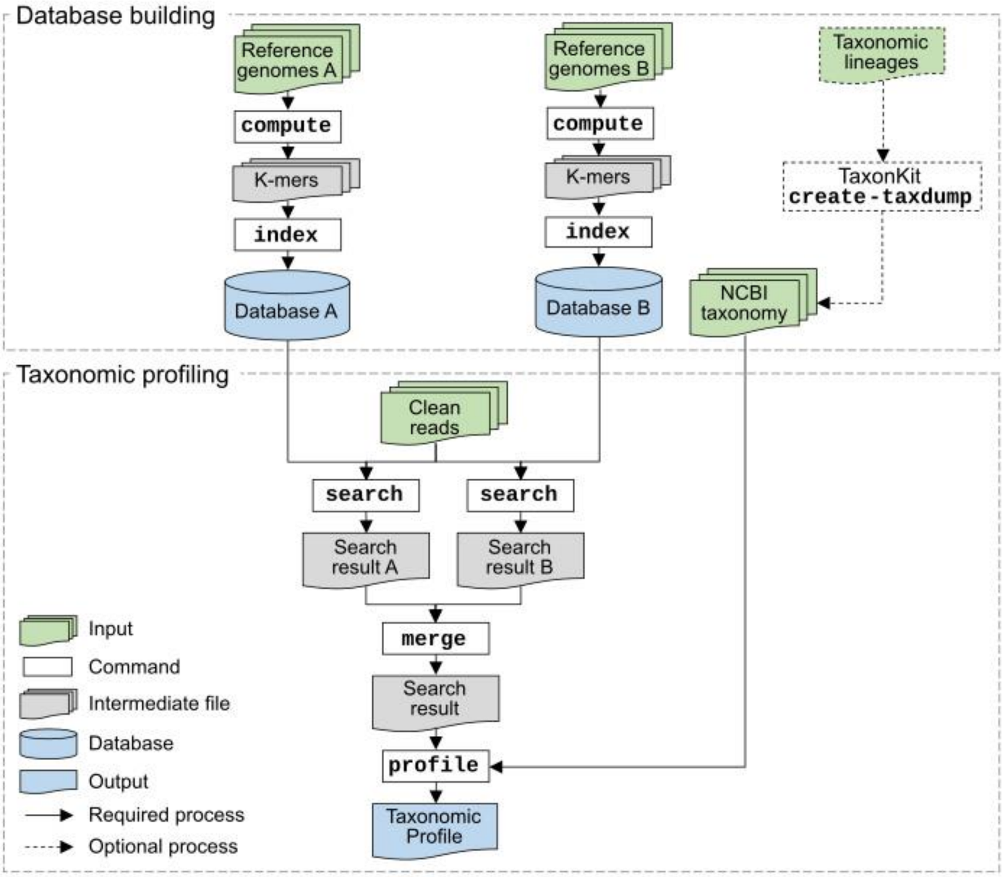

**Figure S1. Workflow of KMCP subcommands**.

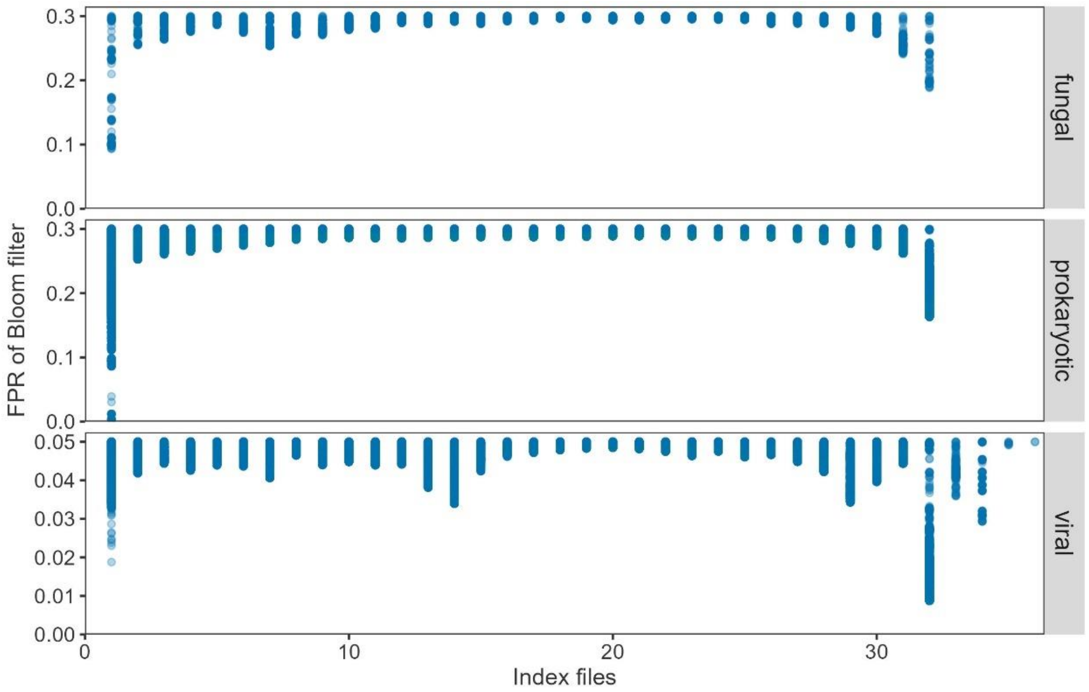

**Figure S2**. **False-positive rates of bloom filters in index files**. The defined false-positive rate of a index block is invalid only for the bloom filter with the most *k*-mers, and other bloom filters have the same or lower false-positive rate due to having less *k*-mers. The defined false-positive rate of the viral database is set to 0.05 in consideration of the smaller genome size.

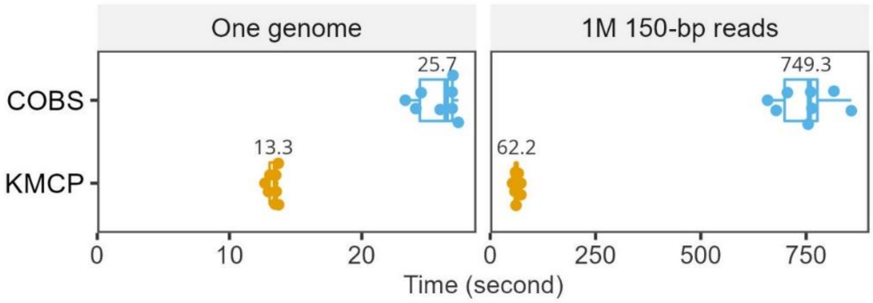

**Figure S3. Benchmarks of searching speed on GTDB dataset**. The 47,894 bacterial and archaeal genomes from GTDB r202 representative collection are used to build databases with the whole *k*-mers ($k = 31$), and eight bacteria genomes (4-5 Mb) are used as queries. The complete genomes (left) or short reads (right) generated from the genomes are used for searching with 40 threads using COBS (commit 1915fc0) and KMCP v0.9.0.

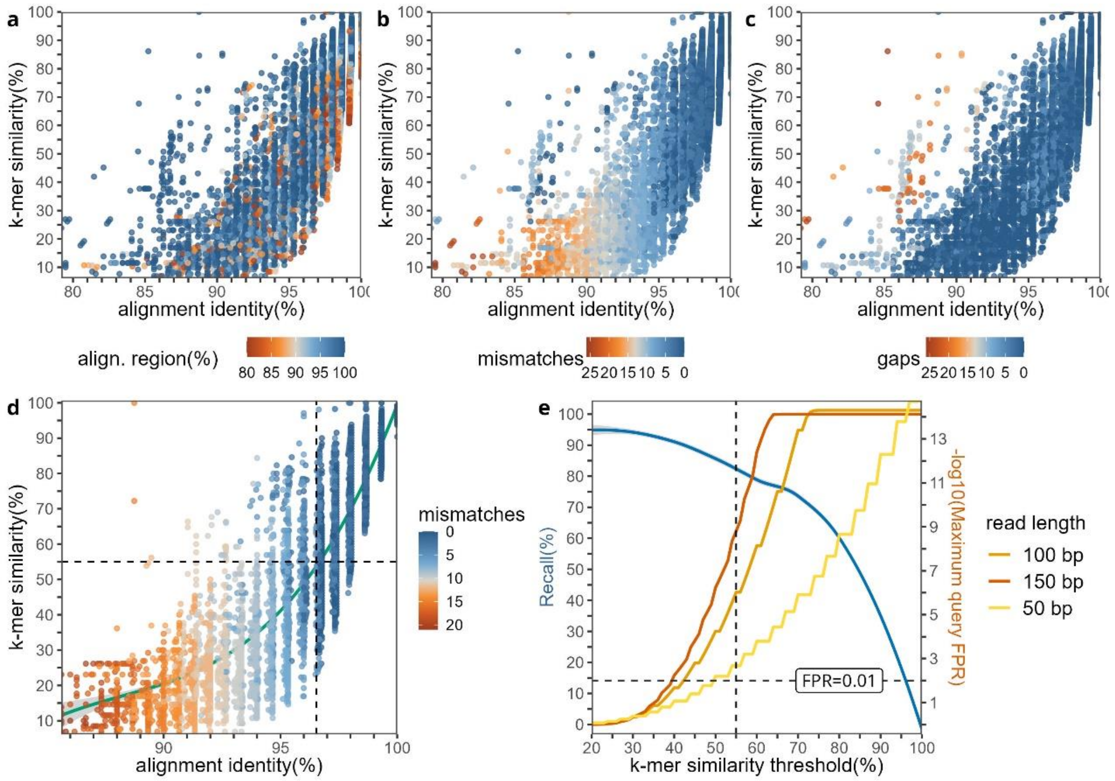

**Figure S4. *K*-mer similarity vs. percent nucleotide identity**. Simulated 150-bp reads are generated from five *Escherichia coli* genomes (GCA_000613265.1, GCA_900706755.1, GCF_000690815.1, GCF_000734955.1, GCF_003697165.2), and aligned to all the genomes with BLASTN. Conserved across all strains but imperfect matches with the percentage of alignment region ≥ 80% are kept, with the best one hit for a search subject. The *k*-mer similarity (percentage of matched *k*-mers for a read) and the alignment identity along with the percentage of the aligned region **(a)**, the number of mismatches **(b)**, and the length of gaps **(c)** are plotted. **(d)** Matches are further filtered with percent identity > 85%, gap length ⩽ 5, and the percentage of aligned regions ≥ 95%. Then a polynomial model of degree 3 is fitted. **(e)** Correlation between *k*-mer similarity threshold and recall of matches and the false-positive rate for reads searching against a KMCP database constructed with one hash function and false-positive rate of 0.3 for each bloom filter.

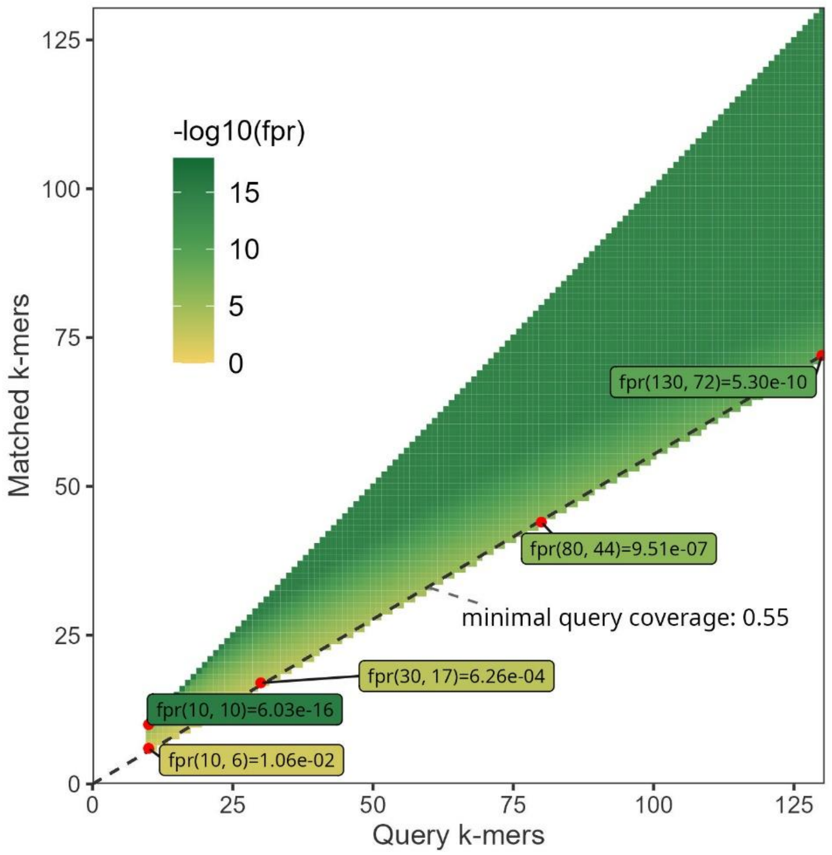

**Figure S5. False-positive rates of queries**. A minimum $k$-mer ($k = 21$) coverage of 0.55 is required for reads ranging from 30 bp to 150 bp. The false-positive rate (FPR) of a single $k$-mer is 0.3, and FPRs of all possible matches are computed according to Theorem 2 in the Solomon and Kingsford paper, which is also implemented in *query-fpr* command in KMCP. For example, a read of 100 bp with 80 (100-21+1) $k$-mers has 44 $k$-mer matched, then the FPR is 9.51e-07.

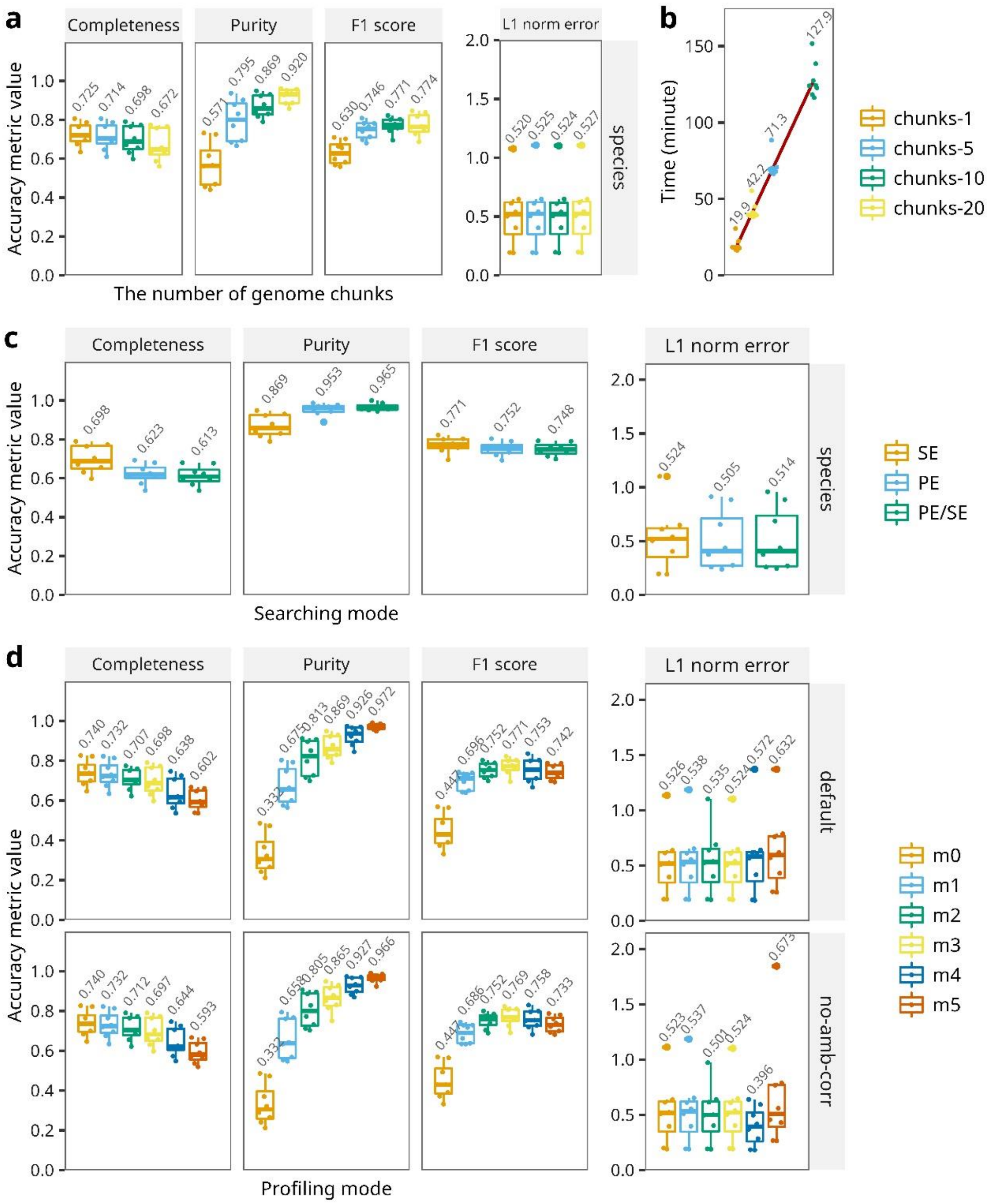

**Figure S6. Accuracies of KMCP on CAMI moust gut dataset with different parameters. (a)** Snapshots of RefSeq and NCBI Taxonomy dump files on Jan 08, 2019, are used to create KMCP database with genome chunks of 1, 5, 10, and 20, respectively. And eight simulated mouse gut datasets are used to perform taxonomic profiling with the four databases respectively. The running time, including the searching and profiling steps, are shown in **(b). (c)** Reference genomes are split into 10 chunks and query reads are searched as single-end (SE), paired-end (PE), or paired-end OR single-end (PE/SE) where PE reads are searched as SE if there are no matches when searching as PE. **(d)** Reference genomes are split into 10 chunks and taxonomic profiling are performed with the six profiling modes respectively. "no-amb-corr" means the flag "—no-amb-corr" is switched on where the two-stage taxonomy assignment algorithm in MegaPath is not used.

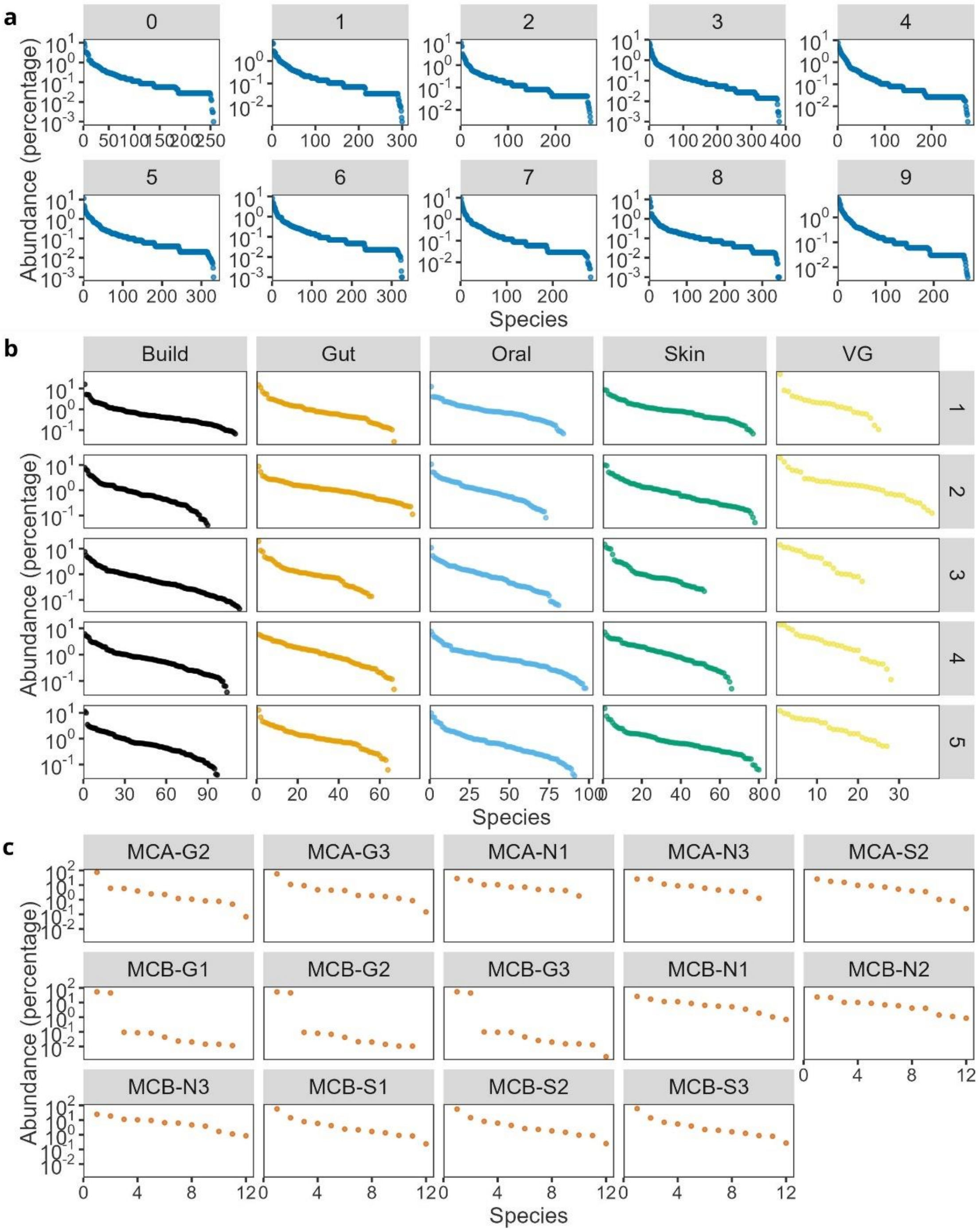

**Figure S7**. **Species abundances in benchmark samples**. Species of the reference genomes are sorted in descending order of percentage of taxonomic abundance. **(a)** The CAMI marine datasets. **(b)** The simulated prokaryotic communities. **(c)** The mock virome datasets.

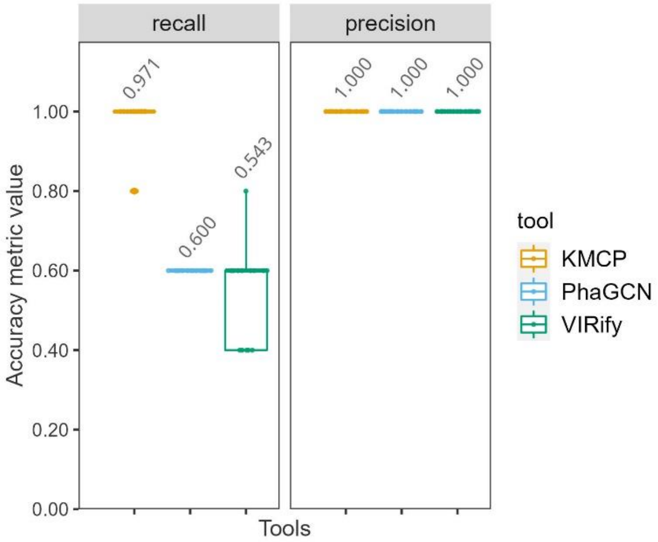

**Figure S8. Taxon identification accuracies at the family rank on the mock virome dataset.** The lowest rank supported is Family for PhaGCN and Genus for VIRify, therefore we compared taxon identification accuracies at the family rank.
